# A brain immune phenotype independent of Alzheimer’s Disease associates with amyloid susceptibility and systemic inflammation

**DOI:** 10.64898/2026.10.21.760450

**Authors:** Giulia Maia Serratto, Gerardo Garcia-Diaz Barriga, Daniel Rosebrock, Malte Simon, Rainer Kuhn, Heinz von der Kammer, Doris Gruber-Schoffnegger

## Abstract

Transcriptomic and proteomic studies have consistently demonstrated substantial molecular heterogeneity across patients with Alzheimer’s disease (AD) and other neurodegenerative disorders, leading to the proposal of multiple disease subtypes. However, almost all previous studies have focused on heterogeneity within diseased cohorts, whereas comparatively little attention has been paid to the molecular diversity of neurologically healthy individuals. In the present study, we demonstrate that one of the most prominent transcriptomic programs characterized by a strong immune component previously interpreted as an AD-specific immune subtype is in fact a brain phenotype that exists independently of AD yet interacts with AD susceptibility and is selectively associated with amyloid pathology. The transcriptional effect of the immune phenotype exceeds that of AD pathology-induced changes and is accompanied by alterations of brain cellular compositions and a distinct systemic inflammation. Our findings suggest that part of the molecular heterogeneity observed in neurological diseases might arise from pre-existing brain phenotypes rather than representing disease-specific subtypes. These insights support a precision medicine approach for Alzheimer’s disease, whereby individuals with an immune phenotype of the brain could be prospectively identified and monitored, enabling the application of targeted preventive strategies before substantial neuropathology and cognitive decline emerge.

## Introduction

Alzheimer’s Disease (AD), the leading cause of dementia, is a clinically and biologically heterogenous neurodegenerative disease characterized by accumulation of extracellular beta-amyloid (Aβ), followed by formation of neurofibrillary tangles (NFT) and progressive neuronal dysfunction (1). Although Aβ deposition initiates 15–20 years before clinical symptoms, NFT accumulation in specific cortical regions more closely tracks with cognitive decline and brain atrophy (2). Yet, individuals with comparable neuropathological burden can exhibit markedly different clinical trajectories, highlighting substantial biological heterogeneity that cannot only be explained by the pathological hallmarks of AD. While rare familial forms of AD are driven by mutations affecting Aβ production and processing, genome-wide association studies (GWAS) of sporadic, late-onset AD have established immune cells as significant determinants of AD risk (3, 4). These findings together with the evidence of inflammatory changes preceding substantial Aβ accumulation, have raised the possibility that alterations in immune homeostasis may contribute to the earliest changes of disease, potentially even before Aβ aggregation (5–8). Persistent or unresolved inflammation could therefore alter brain homeostasis and shape the biological environment in which AD pathology develops in a subset of patients.

The brain is an immunologically active organ in which resident glial cells, including microglia and astrocytes, interact dynamically with vascular and barrier-associated cells as well as peripheral immune signals to maintain tissue homeostasis and shape responses to disease (9). Changes in cell type abundance and cellular transcriptional states are recognized as important features of neurodegenerative diseases but have often been interpreted in the context of disease-associated processes (10–12), without establishing whether they also reflect broader phenotypes that occur independently of disease. Accumulating evidence indicates that systemic inflammation, vascular dysfunction and peripheral immune activation can influence CNS function and inflammatory responses (13–16), but whether such coordinated central and peripheral immune states contribute to the biological heterogeneity observed in AD remains poorly understood. Recent studies of molecular heterogeneity in AD have identified an “immune” subtype, characterized by elevated inflammatory signatures and concurrent neuronal loss (17, 18). Importantly, proteomic analyses discovered that an APOE ε4-associated immune signature was shared across multiple neurodegenerative diseases and cognitively unimpaired individuals, suggesting that at least some immune-related molecular changes reflect broader biological processes rather than AD-specific phenotypes (19). Despite these clues, only limited attention has been given to disentangling disease-specific trajectories and variation in independent phenotypes which shape vulnerability to the disease. Differentiating AD-associated immune activation from inflammation caused by other factors could therefore refine the identification of disease-associated molecular signatures, patient subpopulations, biomarkers and therapeutic targets.

Working within this frame, we first re-analyzed transcriptomic data from the MSBB cohort (18) to determine whether the immune phenotype identified previously in AD is also present in neurologically healthy controls (CTR). We examined its relationship with AD diagnosis, aging, APOE genotype and neuropathological features, extending the analyses across multiple datasets and brain regions. Finally, we integrated brain and peripheral tissue transcriptomic datasets to determine whether a brain immune phenotype is accompanied by coordinated molecular changes in peripheral organs.

Together, these analyses reveal an immune phenotype that is not restricted to AD but occurs in both neurologically healthy and diseased individuals. This phenotype is associated with altered cellular composition, is increasingly prevalent across the aging brain and is enriched in AD patients and APOE ε4 carriers. The regional occurrence of the immune phenotype is differentially associated with Aβ pathology, cognitive outcomes and resilience, while its broader distribution across the brain is accompanied by coordinated inflammatory transcriptional changes in peripheral tissues. These findings suggest that biological processes that precede or occur independently of AD may contribute to inter-individual variation in disease susceptibility and should be considered when defining disease-associated molecular signatures and developing therapeutic strategies.

## Results

### A distinct transcriptomic signature identifies immune-type samples in AD and CTR brains

Transcriptomic analyses have previously highlighted substantial molecular heterogeneity in Alzheimer’s disease and proposed the existence of distinct disease subtypes characterized by specific cellular programs. In particular, Neff et al. (18) identified three macrotypes (A, B and C), further refined into 5 subtypes (A, B1, B2, C1, C2) using parahippocampal gyrus (PHG) samples from AD patients. Interestingly, macrotype C (composed of C1 and C2) was characterized by an elevated immune-associated transcriptional signature and reduced neuronal gene expression. Whether this immune-associated macrotype is specific to AD pathology, however, has not been addressed.

To address this question, we re-analyzed the MSBB PHG transcriptomic dataset by stratifying samples according to neuropathological diagnosis (AD only, CTR only, or all samples combined). To define clusters, we applied Weighted Sample Correlation Network Analysis (WSCNA), an unbiased clustering algorithm that groups transcriptionally similar samples using pairwise expression correlations. We identified three macroclusters independent of the stratification applied, that could be further refined into 5 clusters in AD and in combined samples (Fig. S1, A-C). Gene signatures of each cluster were highly consistent with previously defined subtypes and sample annotation could be robustly replicated (Fig. S1, D-I). The existence of the previously defined macrotype C in control samples indicates that the immune signature is not specific to AD pathology and does not describe a brain subtype exclusive to Alzheimer’s diagnosis. Therefore, to establish a consistent terminology throughout this study, we selected the combined-cohort clusters as our reference, retaining the original annotations (A, B1, B2, C1, C2). We defined “immune type” the sample classification for macrocluster C, whereas macroclusters A and B are collectively designated as “other type”. We refer to the underlying gene expression pattern as the “immune signature” and use “immune phenotype” to describe the broader biological trait characterized by this classification.

To prove that the immune-type classification of samples is independent of diagnosis, we applied WSCNA clustering to temporal cortex (TC) samples from AD patients and CTRs collected from three different datasets (MSBB, ROSMAP, MAYO). Upon correction for sex, study and batch, we identified three clusters having high similarity to the reference clusters (Fig. 1, A-F), with cluster C (purple) identified as the corresponding immune type. All clusters could be detected in both AD and CTR samples (Fig. 1G).

**Fig. 1.**
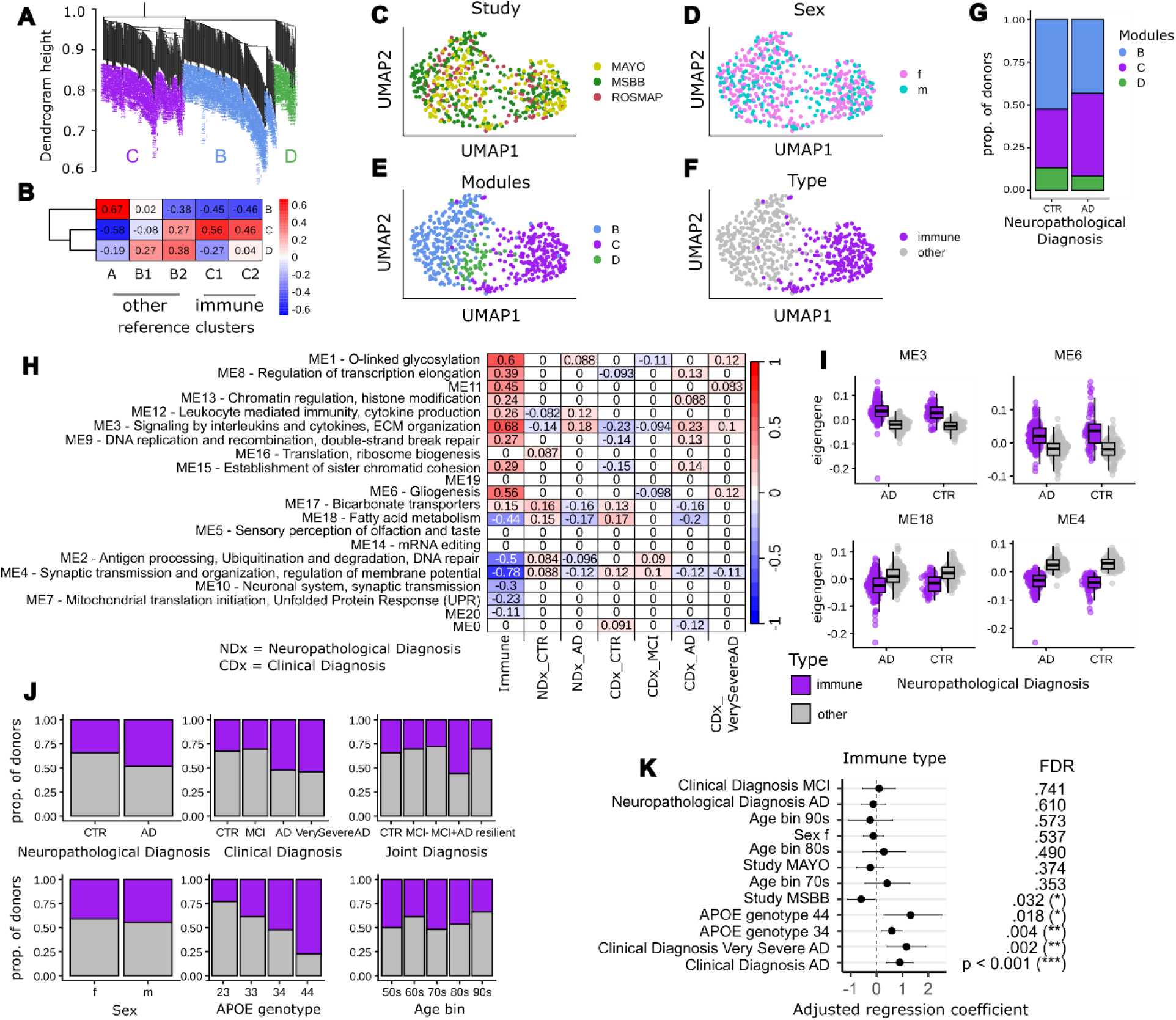
Identification and characterization of a conserved transcriptomic immune type across independent AD and control cohorts. **(A)** Weighted Sample Gene Network Analysis (WSCNA) clustering dendrogram of temporal cortex (TC) samples merged from the MSBB, ROSMAP, and MAYO cohorts, resolving into three distinct sample clusters: B (blue), C (purple), and D (green). **(B)** Pairwise similarity heatmap based on a modified Jaccard index comparing the marker gene signatures of the newly identified TC clusters (B, C, D) to the original reference macro-types and subtypes (A, B1, B2, C1, C2). Cluster C exhibits high transcriptomic concordance with reference clusters C1 and C2, defining it as the "immune type." **(C-F)** Uniform Manifold Approximation and Projection (UMAP) embeddings of the TC samples, color-coded by study cohort (C), donor sex (D), identified WSCNA modules (E), and assigned reference macro-type (F). **(G)** Relative donor distribution across clusters B, C, and D, stratified by neuropathological diagnosis. **(H)** Correlation matrix between Weighted Gene Co-expression Network Analysis (WGCNA) gene modules (rows) and clinical (CDx) or pathological (NDx) diagnosis (columns). Cell values report the correlation coefficients for statistically significant associations (p < 0.05); non-significant values are indicated by 0. Row names highlight the module identifier and its top enriched biological pathways (comprehensive pathway enrichments are detailed in Table S1). **(I)** Boxplots displaying WGCNA module eigengenes for four representative modules (ME3, ME6, ME18, ME4) strongly correlated with the immune phenotype, separated by type assignment (immune vs. other) and neuropathological diagnosis. **(J)** Relative proportions of the immune type across sample stratifications, including neuropathological, clinical, and joint diagnosis, donor sex, APOE genotype, and age bins. **(K)** Forest plot illustrating the adjusted regression coefficients (points) and corresponding 95% confidence intervals (error bars) from a generalized linear model identifying predictors of the immune phenotype. Models are simultaneously adjusted for study, age bin, sex, APOE genotype, and clinical/neuropathological status. Dashed line at 0 indicates the null hypothesis. Asterisks denote statistical significance: *FDR < 0.05, **FDR < 0.01, ***FDR < 0.001. NDx, Neuropathological Diagnosis; CDx, Clinical Diagnosis; CTR, Control; MCI, Mild Cognitive Impairment; AD = Alzheimer’s Disease.

We next performed Weighted Gene Coexpression Network Analysis (WGCNA), – an unsupervised method that identifies highly correlated gene modules and links them to specific phenotypic traits – to further characterize the immune type (Fig. 1H, Table S1). The module with the greatest positive correlation with the immune-type classification was enriched for pathways including innate immune signaling, antiviral responses, and extracellular matrix organization (ME3; r = 0.68) with *MAVS* and *TRIM56,* pivotal regulators of viral immune signaling, as top associated genes (Table S1). Other top-ranked modules tracking positively with the immune type were enriched for O-linked glycosylation (ME1; r = 0.60) and gliogenesis (ME6; r = 0.56). Conversely, the most strongly down-regulated modules were enriched for synaptic and neuronal pathways (ME4; r = -0.78), featuring key hub genes including the growth factor *FGF12*, calcium-dependent calmodulin-stimulated *PPP3CA* and *PPP3CB*, and the GABA receptor subunits *GABRA1* and *GABRB2*. Other top down-regulated modules were enriched for ubiquitination/degradation (ME2; r = -0.50) and fatty acid metabolism (ME18; r = -0.44). While a broader suite of modules reached statistical significance (e.g., ME8, ME12, ME9), they exhibited notably weaker effect sizes (r < 0.4). Interestingly, only a subset of modules showed weak associations with AD diagnosis, reinforcing the notion that the immune signature dominates over AD-driven transcriptomic changes. Therefore, the immune-type classification captured biological variation that was not solely explained by AD diagnosis. To directly demonstrate that these co-expression dynamics are inherently driven by the immune type rather than by diagnosis, we examined the eigengenes of 4 top correlating modules (ME3/4/6/18) across immune-type and other samples (Fig. 1I). Eigengene values, representing the first principal component that captures the summary expression profile of a module, were driven by cluster identity rather than diagnosis. In fact, the upregulation of ME3 and ME6 and the downregulation of ME4 and ME18 in the immune-type samples were independent of neuropathological status. We next evaluated if the immune type was overrepresented in samples from donors with AD diagnosis or if it was associated with age, sex or APOE genotype (Fig. 1J, K). The relative abundance of the immune type was significantly higher in donors clinically diagnosed with AD compared to controls, but not in those with MCI, and was significantly associated with APOE ε4 genotype. No significant associations were seen with age or sex. These observations indicate that the immune type constitutes a distinct transcriptional state that can occur independently of neuropathological status, yet may influence susceptibility to AD, as evidenced by its increased prevalence among AD brain samples.

### Regional immune-type classification differentially associates with AD pathology and cognitive impairment

As initial analyses focused on evaluating immune-type samples in a single brain region, we next investigated whether multiple areas of the same brain could be consistently assigned to the immune type. We leveraged multi-region bulk transcriptomic data from ROSMAP, which includes up to 5 brain regions per donor - dorsolateral prefrontal cortex (DLPFC), frontal cortex (FC), temporal cortex (TC), posterior cingulate cortex (PCC), head of caudate nucleus (HCN). We applied WSCNA and identified 5 transcriptionally distinct clusters after correction for sex, brain region and batch (Fig. 2, A-C). Based on similarity to the reference set, clusters C and E collectively constitute the immune type in the ROSMAP cohort (Fig. 2, D and E). Notably, the same ROSMAP TC samples were used in the multi- cohort analysis (Fig. 1) and the comparison of the two independent rounds of clustering revealed robust sample annotation (Table S2) confirming that pooling multiple brain regions does not alter individual cluster identities. To assess the spatial consistency of the immune type within the brain, we quantified the proportion of immune-type samples for donors with at least 4 brain regions available (n = 110, Fig. S2) and called this parameter “prop_immune”. While 30% of donors demonstrated high consistency of the immune type across the brain (prop_immune ≥ 0.75), 36% displayed variable regional assignment (0 < prop_immune < 0.75), and 34% showed no immune-type regions (prop_immune = 0) (Fig. 2F). Dichotomizing prop_immune into low (prop_immune < 0.25) and moderate-to-high (prop_immune ≥ 0.25) groups, highlighted a significant enrichment of moderate-to-high prop_immune in donors diagnosed with AD (clinical and joint diagnosis; definitions in Methods), and a trend towards a gene dosage-dependent increase in APOE ε4 carriers, though this did not reach significance when comparing APOE ε4 carriers directly against non-carriers (Fig. 2G). These observations suggest that while regional assignment to the immune type does not necessarily reflect a brain-wide phenotype, it nonetheless associates with AD diagnosis.

**Fig. 2.**
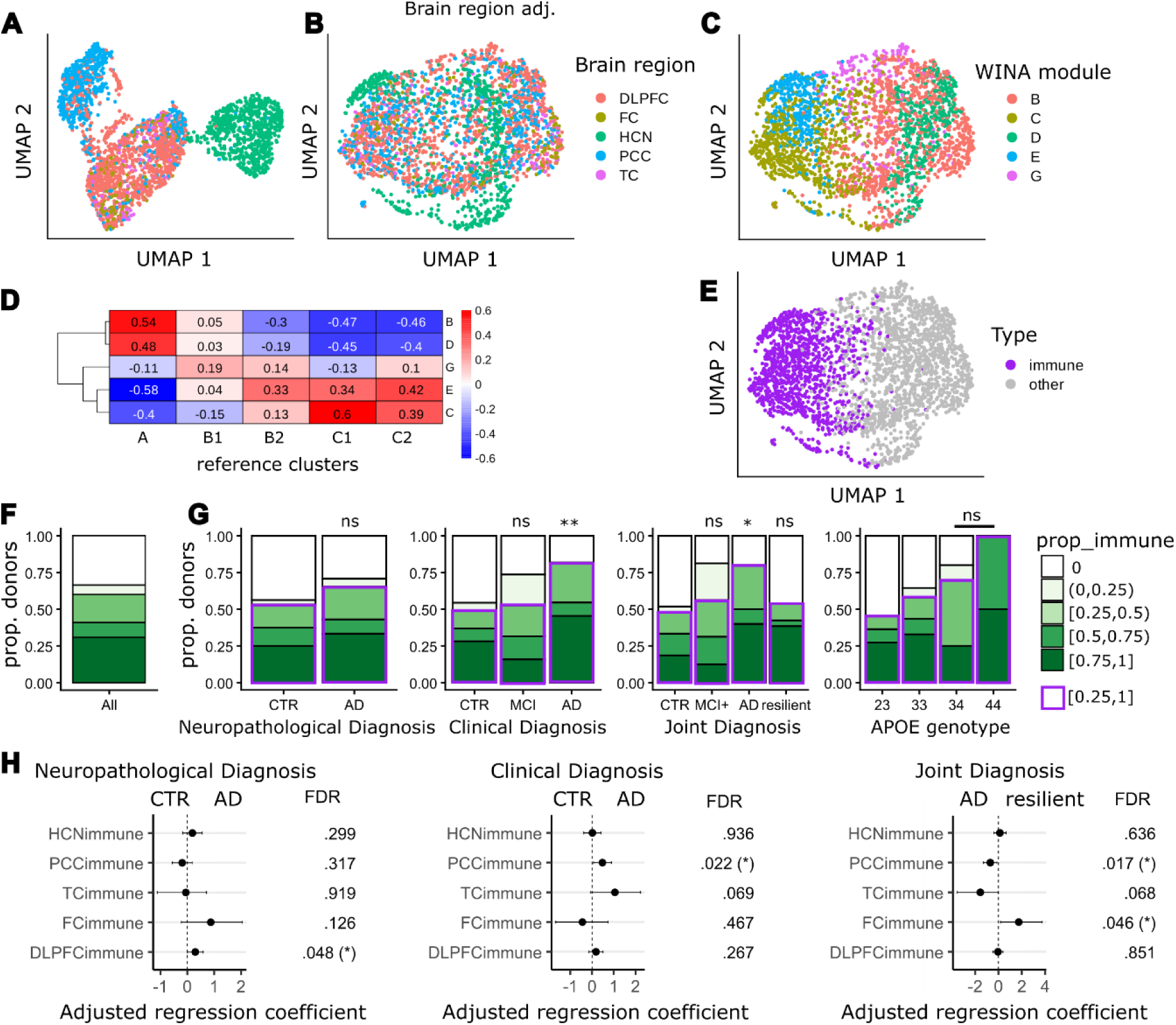
Regional immune-type classification reveals distinct associations with AD pathology and cognitive impairment. **(A and B)** UMAP embeddings of ROSMAP multi-region bulk transcriptomic samples colored by brain region before (A) and after (B) batch-effect correction and regression of regional baseline signatures. **(C)** UMAP visualization of samples annotated by the five transcriptionally distinct clusters identified via the WSCNA framework. **(D)** Modified Jaccard index heatmap validating the transcriptomic overlap between the five multi-region clusters and the established reference signatures. Clusters C and E correlate with reference clusters C1 and C2, respectively, collectively representing the immune type within the ROSMAP cohort. **(E)** UMAP visualization of sample clustering. Each dot represents one sample. Samples classified as immune type are highlighted in purple, while all other samples are shown in gray. **(F)** Overall donor distribution of prop_immune (the proportion of immune- type samples per individual) restricted to donors with ≥ 4 evaluated brain regions. **(G)** Donor distribution of prop_immune stratified by neuropathological, clinical, and joint diagnoses, alongside *APOE* genotype. Purple borders highlight fractions of individuals with prop_immune ≥ 0.25. Statistical significance (showed above bars) is derived from a quasi-binomial logistic regression model adjusting for sex, where all diagnosis groups are compared against CTR, and *APOE* genotype comparisons evaluate *APOE ε4* carriers versus non-carriers. * p < 0.05, ** p < 0.01, ns = non-significant. **(H)** Forest plots showing adjusted regression coefficients and 95% confidence intervals from generalized linear models assessing regional immune-type classifications as predictors of neuropathological, clinical, or joint AD diagnoses. DLPFC, dorsolateral prefrontal cortex; TC, temporal cortex; HCN, head of caudate nucleus; FC, frontal cortex; PCC, posterior cingulate cortex.

We next examined whether an immune-type classification in specific brain regions was associated with AD outcomes. Immune-type DLPFC samples were significantly associated with neuropathological diagnosis (overall burden of Aβ and NFT), whereas immune-type PCC samples were significantly associated with clinical AD diagnosis (cognitive decline) (Fig. 2H). Notably, immune-type PCC samples were significantly associated with reduced odds of cognitive resilience in the presence of high pathological burden, reinforcing the importance of the transcriptional profile of this area for cognition. Immune-type FC samples were associated with AD pathological burden, while immune-type TC samples were associated with clinical cognitive decline, though neither association reached statistical significance.

Overall, these observations indicate that the consistency of the immune type across brain regions varies substantially between individuals, and that its association with AD pathology or cognitive decline is region dependent.

### The immune-type DLPFC selectively associates with Aβ deposition

Given the significant association between the DLPFC immune type and AD neuropathology, we next investigated whether this relationship was more closely linked to Aβ deposition or NFT burden. While overall neocortical Aβ levels were greater in AD compared to CTR, they were significantly elevated in donors with an immune-type DLPFC within both neuropathological diagnosis and CERAD neuritic plaque score stratifications (Fig. 3A, B). Across the age continuum, AD donors with an immune-type DLPFC consistently exhibited higher neocortical Aβ levels compared to those with an other-type DLPFC (Fig. 3C). In contrast, among CTR donors, Aβ levels increased with age exclusively in individuals with an immune-type DLPFC. No significant differences in NFT burden were observed between donors with or without immune-type DLPFCs (Fig. 3D, E). Immune-type classification of the DLPFC did not modify the relationship between NFT burden and age in either AD or CTR donors (Fig. 3F). Despite the elevated Aβ burden, longitudinal rates of cognitive decline did not differ significantly between the two groups, including when samples were stratified by Braak stage (Fig. 3G). Together, these findings link the immune-type DLPFC more specifically to Aβ deposition than to NFT burden, without evidence of an association with accelerated cognitive decline.

**Fig. 3.**
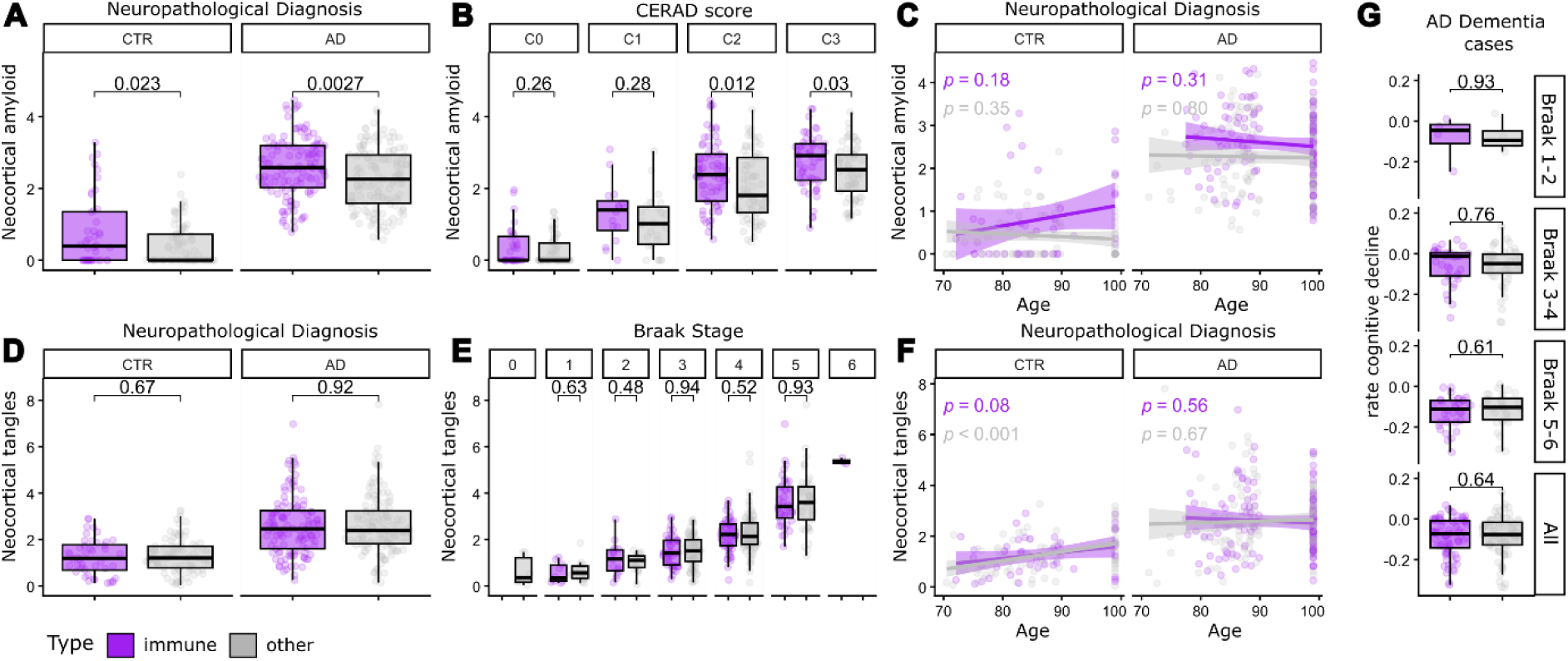
Divergent associations of the immune-type DLPFC with amyloid versus tau pathology **(A and B)** Boxplots comparing global neocortical Aβ deposition between individuals exhibiting an immune-type (purple) vs. other-type (grey) DLPFC, stratified by overall neuropathological diagnosis (A) or CERAD neuritic plaque score (B). **(C)** Scatter plot displaying the correlation between neocortical Aβ burden and age, colored by DLPFC sample classification and stratified by neuropathological diagnosis. Correlation p-value is displayed on top. **(D and E)** Boxplots comparing neocortical NFT burden between immune-type and other-type DLPFC samples across overall neuropathological diagnoses (D) and Braak stages (E). **(F)** Scatter plot highlighting the correlation between neocortical NFT burden and age, colored by DLPFC classification and stratified by neuropathological diagnosis. **(G)** Boxplots comparing the rate of cognitive decline across progressive Braak stages in clinically diagnosed AD dementia cases, between donors with and without an immune-type DLPFC. Pairwise comparisons between immune-type and other samples are computed using two-tailed Student’s t-tests. Correlation significance was evaluated using a two-tailed Pearson correlation test. Exact p-values are reported directly above brackets or within the plot.

### The immune-type DLPFC exhibits distinct cellular composition and specific response to AD pathology

Because the immune signature is enriched in immune and neuronal genes, immune-type samples might be characterized by an altered cellular composition. Therefore, we compared the relative abundance of the major brain cell types between immune-type and other DLPFC samples. For this analysis, we utilized the cellular annotations recently published by Green et al. (20) for the subset of ROSMAP DLPFC samples with available snRNA-Seq data (n = 421). After adjusting for sex, age and neuropathological diagnosis, immune-type DLPFC samples displayed a significant shift in cellular composition, characterized by decreased proportions of both inhibitory and excitatory neurons and a corresponding increase in astrocyte, microglia, pericyte, endothelial cell and OPC abundance (Fig. 4A). We next investigated whether AD neuropathology evoked specific changes in immune-type or other DLPFC samples. While comparisons between AD and CTR samples revealed a significant reduction in inhibitory neurons and an increase of oligodendrocytes in AD, stratification by transcriptomic type resulted in no significant difference between AD and CTR samples.

**Fig. 4.**
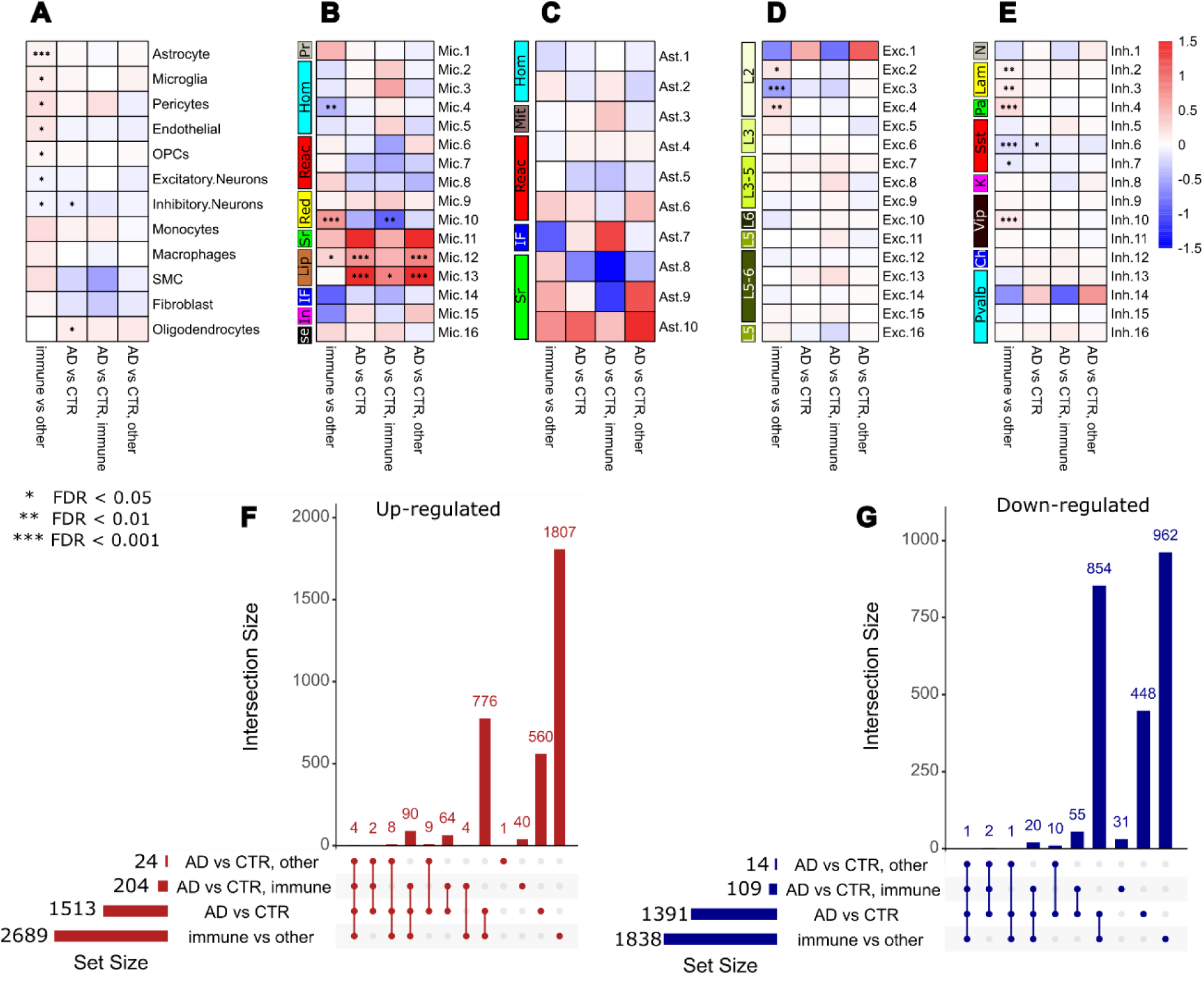
The immune-type DLPFC exhibits distinct cellular composition and specific response to AD pathology **(A-E)** Heatmaps depicting comparative changes in the relative abundance of major central nervous system (CNS) cell types (A) and fine-grained subpopulations of microglia (B), astrocytes (C), excitatory neurons (D), and inhibitory neurons (E). Proportions are derived from ROSMAP DLPFC single-nucleus RNA sequencing (snRNA-Seq) data. Values reflect log-transformed effect sizes from quasi-binomial regression models accounting for key confounding covariates across four distinct comparison axes: immune type vs. other-type, AD vs. CTR, AD vs. CTR restricted to immune-type samples, and AD vs. CTR restricted to other-type samples. Asterisks indicate Benjamini-Hochberg false discovery rates: *FDR< 0.05, **FDR< 0.01, ***FDR < 0.001. In B and C: Pr, proliferating; Hom, homeostatic; Reac, reactive; Red, Enhanced redox; Sr, stress response; Lip, lipid associated; IF, interferon response; In, inflammatory; Se, SERPINE1+; Mit, enhanced mitophagy; in D: L, cortical layer; in E: N, NPY+; Lam, LAMP5+; Pa, PAX6+; Sst, SST+; K, KIT+; Vip, VIP+; Ch, Chandelier; Pvalb, PVALB+. **(F and G)** UpSet plots detailing the intersections, unique components, and overall set sizes of significantly upregulated (F) and downregulated (G) differentially expressed genes (DEGs) computed from ROSMAP bulk transcriptomic profiles across the corresponding four comparison axes. Stringency thresholds were applied to define DEGs (|log2FC|> 0.1, FDR < 0.05 for pathology axes; |log2FC|> 0.5, FDR < 0.05 for the immune type axis).

To determine whether the altered cellular composition of immune-type DLPFC samples was accompanied by changes in specific cellular subpopulations, we compared the relative abundance of microglial, astrocytic, and neuronal subtypes. Among microglia (Fig. 4B), the homeostatic Mic.4 population was significantly depleted in immune-type DLPFC samples, whereas TMEM163⁺ Mic.10 microglia, characterized by endoplasmic reticulum (ER) stress, ribosomal gene expression, and inflammatory markers, were significantly enriched. The lipid-associated CPM⁺ Mic.12 population, previously linked primarily to Aβ pathology (20), was also significantly enriched in immune-type DLPFC.

Comparison between AD and CTR samples further revealed increased abundance of the lipid-associated microglia Mic.12 and PTPRG^+^ Mic.13 populations, the latter previously implicated as a potential causal contributor to tau pathology. Both populations remained significantly enriched in AD when the analysis was restricted to other-type samples. Within immune-type DLPFC samples, however, AD was associated with a reduction in Mic.10 abundance and an increase in PTPRG^+^ Mic.13 relative to controls, further supporting the association of this microglial population with AD pathology.

Collectively, these observations suggest that the increased abundance of microglia in immune-type DLPFC is accompanied by a depletion of homeostatic and an enrichment of distinct reactive populations. The data also suggests that the relationship between AD pathology and specific microglial subpopulations varies according to the transcriptomic classification of the tissue.

Although astrocytes showed the strongest enrichment in the immune-type DLPFC, there was no change in the relative abundance of any astrocyte subpopulations (Fig. 4C). Furthermore, AD samples did not show any significant alterations in astrocyte subpopulations compared to CTR.

Among excitatory neurons, layer 2 (L2) neurons were selectively affected, showing a significant reduction in Exc.3 abundance and concomitant increase in Exc.2 and Exc.4 in immune-type compared to other-type DLPFC samples (Fig. 4D). Among inhibitory neurons, the SST^+^ interneuron subpopulations Inh.6 and Inh.7, were significantly depleted in immune-type DLPFC, whereas VIP^+^ Inh.10, PAX6^+^ Inh.4, LAMP5^+^ Inh.2 and Inh.3 were significantly increased (Fig. 4E).

When comparing AD samples to CTR irrespective of the transcriptional type, a significant reduction in the SST^+^ Inh.6 subpopulation was observed, while there was no change observed in any neuronal subpopulation when samples were stratified according to immune-type classification.

To compare the transcriptional impact of the immune type and AD pathology on DLPFC samples, we calculated differentially expressed genes (DEGs) between immune-type and other samples (adjusted for sex, age, neuropathological diagnosis), as well as between AD and CTR samples (adjusted for sex, age). The immune signature in the DLPFC was defined by 2689 up- and 1838 down-regulated genes, whereas the AD signature was composed of 1513 up- and 1391 down-regulated genes. (Fig. 4F, G). Interestingly, ∼70% of the AD signature overlapped with the immune signature, highlighting that many gene expression changes conventionally attributed to AD pathology might instead be confounded by or mediated through the immune-type classification. Furthermore, AD signatures calculated within immune-type or other samples were defined by a drastically lower number of genes, despite the high number of samples (immune type: 118 AD, 44 CTR; other type: 136 AD, 88 CTR, Table S2), and showed minimal overlap between them. Only 9 DEGs were significantly altered by AD pathology regardless of sample stratification: *RRAD*, *A2ML1*, *AQP6*, *DIO3*, *GAREM2*, *SPACA6* were up- regulated, *NUDT7*, *SST*, *LOC101927879* were down-regulated (Table S1).

These observations indicate that while a core AD signature exists, most changes associated with AD neuropathology are influenced by the background immune phenotype.

### Brain-wide expansion of the immune phenotype follows major neuronal pathways

Since the immune-type classification of cortical samples is not specific to AD, but is associated with Aβ accumulation, a major hallmark of the disease, the emergence of this phenotype in the cortex may represent a risk for AD development. Therefore, we investigated the brains of younger, neurologically healthy individuals to reveal potential patterns associated with acquiring an immune-type cortex. We leveraged bulk transcriptomic data from the GTEx consortium, with samples from neurologically healthy donors, ranging from 20–79 years of age. The GTEx collection comprises up to 13 brain regions per individual, enabling a comprehensive assessment of variations in the occurrence of the immune- type classification throughout the brain. We applied WSCNA clustering to the entire GTEx brain dataset, adjusting for brain region and sex, and identified 7 transcriptionally distinct clusters (Fig. 5, A- C). Clusters B, G and I showed high similarity to reference clusters C1 and C2 and collectively constituted the immune type in the GTEx cohort (Fig. 5, D and E). We next quantified prop_immune within each donor with at least 10 brain regions (n = 113, Fig. S2). While 14% of donors demonstrated high consistency of the immune-type classification across the brain (prop_immune ≥ 0.75), 66% displayed variable regional assignment (0 < prop_immune < 0.75), and 20% showed no immune-type region (prop_immune = 0) (Fig. 5F). Stratification of donors by age groups highlighted a clear age- dependent increase in prop_immune, suggesting that a highly consistent immune-type classification across the brain is not a static brain characteristic, but might result from the progressive expansion of the immune phenotype across brain regions.

**Fig. 5.**
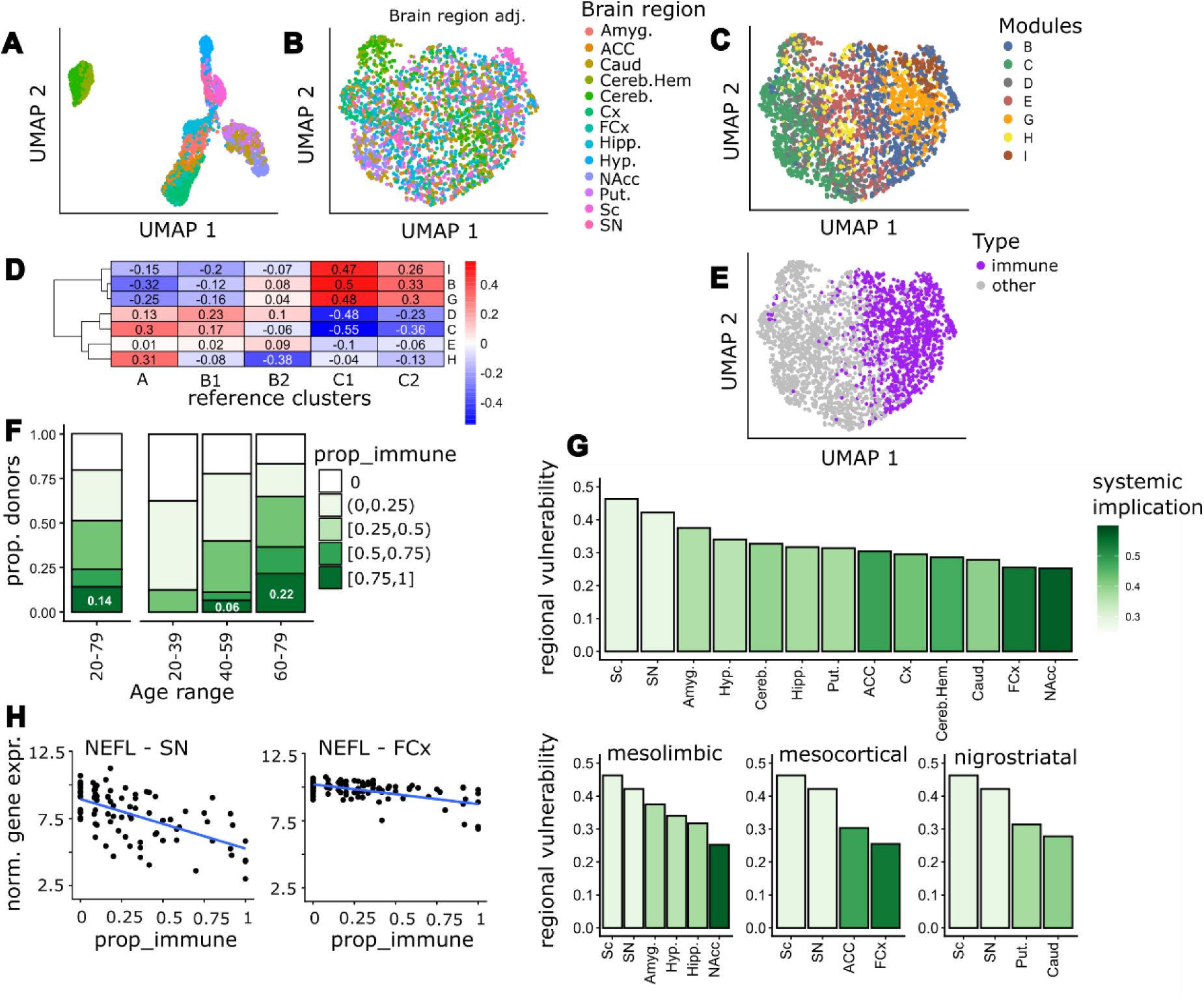
Brain-wide topographical expansion of the immune-type classification follows major neuronal pathways **(A-B)** UMAP representation of transcriptomic profiles from GTEx brain samples (cognitively normal donors, age range 20 – 79 years), color-coded by specific brain areas before (A) and after (B) regressing brain region-specific baseline expression features. **(C)** UMAP colored by WSCNA module assignment. Seven transcriptionally distinct modules (B–I) were identified across the GTEx brain dataset. **(D)** Modified Jaccard index map comparing GTEx clusters with the reference clusters defined in the discovery cohort. Clusters B, G, and I exhibited the highest similarity to the reference C1 and C2 clusters, and were therefore classified as immune type. **(E)** UMAP visualization of immune-type classification. Samples classified as immune type are shown in purple, while all remaining samples are shown in gray. **(F)** Distribution of the proportion of immune-type samples per individual (prop_immune) across age groups. Donors with ≥ 10 regions were included in the analysis. **(G)** Ranked bar chart showing regional vulnerability (the proportion of samples classified as immune type across individual brain regions). Bars are colored by systemic implication (for each specific brain region, the proportion of donors with an immune-type classification in that region who also exhibited a highly consistent immune-type classification across the brain (prop_immune ≥ 0.75)). Upper plot shows all profiled regions, while plots at the bottom stratify regions by three distinct neural pathways. **(H)** Scatter plots demonstrating the differential correlation of a standard neuronal marker (*NEFL*) with prop_immune values in a ventral region (SN) compared to a neocortical region (FCx). Gene expression values are adjusted for age and sex; blue lines show linear regression slopes. Amyg, Amygdala; ACC, Anterior Cingulate Cortex; Caud, Caudate Nucleus; Cereb.Hem, Cerebellar Hemisphere; Cereb, Cerebellum; Cx, Cortex; FCx, Frontal Cortex (BA9); Hipp, Hippocampus; Hyp, Hypothalamus; NAcc, Nucleus Accumbens; Put, Putamen; Sc, Cervical C1 Spinal Cord; SN, Substantia Nigra.

To track the potential routes of expansion of the immune phenotype, we calculated two complementary parameters: 1) regional vulnerability: the proportion of samples classified as immune type within each brain region examined; 2) systemic implication: for each specific brain region, the proportion of donors with an immune-type classification in that region who also exhibited a highly consistent immune-type classification across the brain (prop_immune ≥ 0.75). Joint visualization of the two parameters revealed a striking inverse relationship (Fig. 5, G): brain regions exhibiting the highest proportion of immune-type samples (most vulnerable regions) were the least predictive of a highly consistent immune-type classification across the brain (lowest systemic implication). The most vulnerable regions were the spinal cord (46% of samples classified as immune type) and midbrain/subcortical areas (substantia nigra (SN) and amygdala (Amyg.), 42% and 38% of samples classified as immune type, respectively), while the frontal cortex (FCx) and the nucleus accumbens (NAcc) were the least vulnerable (26% and 25% of samples classified as immune type, respectively). To further characterize the potential route of expansion starting from the most vulnerable region (spinal cord), we grouped the remaining profiled brain areas into three major neural pathways: mesolimbic, mesocortical and nigrostriatal pathways. The SN was included in the mesolimbic and mesocortical pathways assuming samples might also contain the ventral tegmental area, due to the anatomical proximity. The combined analysis of regional vulnerability and systemic implication revealed a coherent spatial gradient that suggested the mesolimbic and mesocortical pathways as major axes of expansion of the immune phenotype. Consistent with this model, the pathway termini, NAcc and the FCx emerged as the regions most predictive of a highly consistent immune-type classification across the brain, with 57% and 54% of donors, respectively, exhibiting prop_immune ≥ 0.75 when the NAcc or FCx had the immune-type classification.

The ordered topographical expansion of the immune phenotype also resulted in a progressive exacerbation of the immune signature in the most vulnerable regions. In fact, expression of NEFL (Fig. 5H) and several neuronal and immune-related genes (Table S1) increased/declined more steeply with respect to prop_immune in the SN than in the FCx, the respective start- and endpoints in the mesocortical pathway.

To capture the effects of the topographical expansion of the immune phenotype on cellular composition, we applied the CelMod deconvolution method to the GTEx FCx samples (Methods, Fig. S3, Fig. S4) and calculated the changes in cell type abundance with increasing prop_immune (Fig. S5, A-E, Table S1). While the relative abundance of both excitatory (r = -0.62) and inhibitory neurons (r = -0.55) significantly decreased with increasing prop_immune, astrocytes (r = 0.62), monocytes (r = 0.54), OPCs (r = 0.52) and vascular cells (endothelial r = 0.73, pericytes r = 0.68, fibroblasts r = 0.37) increased with increasing prop_immune. The relative abundance of microglia (r = 0.11), macrophages (r = 0.27) and oligodendrocytes (r = -0.01) remained unchanged. These alterations partially reflected the changes observed in immune-type DLPFC samples in the snRNA-Seq ROSMAP dataset (Fig. 4). The key distinction between the cellular composition of immune-type samples and the changes associated with its brain-wide distribution concerned microglia. Although microglial abundance was increased in immune-type relative to other samples, it did not vary with prop_immune. This suggests that the broader distribution of the immune type across the brain is associated with dynamic transitions in microglial states rather than further changes in their overall abundance.

Overall, these observations support a model of progressive topographical expansion of the immune type along the mesolimbic and mesocortical pathways. The immune phenotype appears to originate in the spinal cord or more ventral brain structures before gradually reaching the cortex. Within the cortex, the immune signature and the cellular composition of immune-type samples are progressively exacerbated by the expansion of the immune phenotype across the brain.

### Brain-wide expansion of the immune phenotype is linked to coordinated transcriptomic changes in peripheral tissues

To investigate the association between brain-wide expansion of the immune phenotype and molecular changes in peripheral organs, we examined the relationship between increasing prop_immune and gene expression levels in colon, heart and whole blood samples of the GTEx collection (Fig. 6A). RNA-Seq data from colon samples were available for 53 donors (sigmoid n = 44 samples, transverse n = 23 samples) for whom a corresponding brain prop_immune value had also been determined. After adjusting for age, sex, and colon sub-region, we identified 499 genes significantly correlating with prop_immune and 682 showing inverse correlation (Fig. 6, B and C, Table S1). While *ALOX15B,* a marker of gastrointestinal inflammation, exhibited strong correlation with prop_immune and a steep positive slope (slope = 1.4), *FAM69C*, a kinase exerting a protective role against cellular stress, showed one of the most negative values (slope = -1.2) and significant inverse correlation (Fig. 6, D and E).

**Fig. 6.**
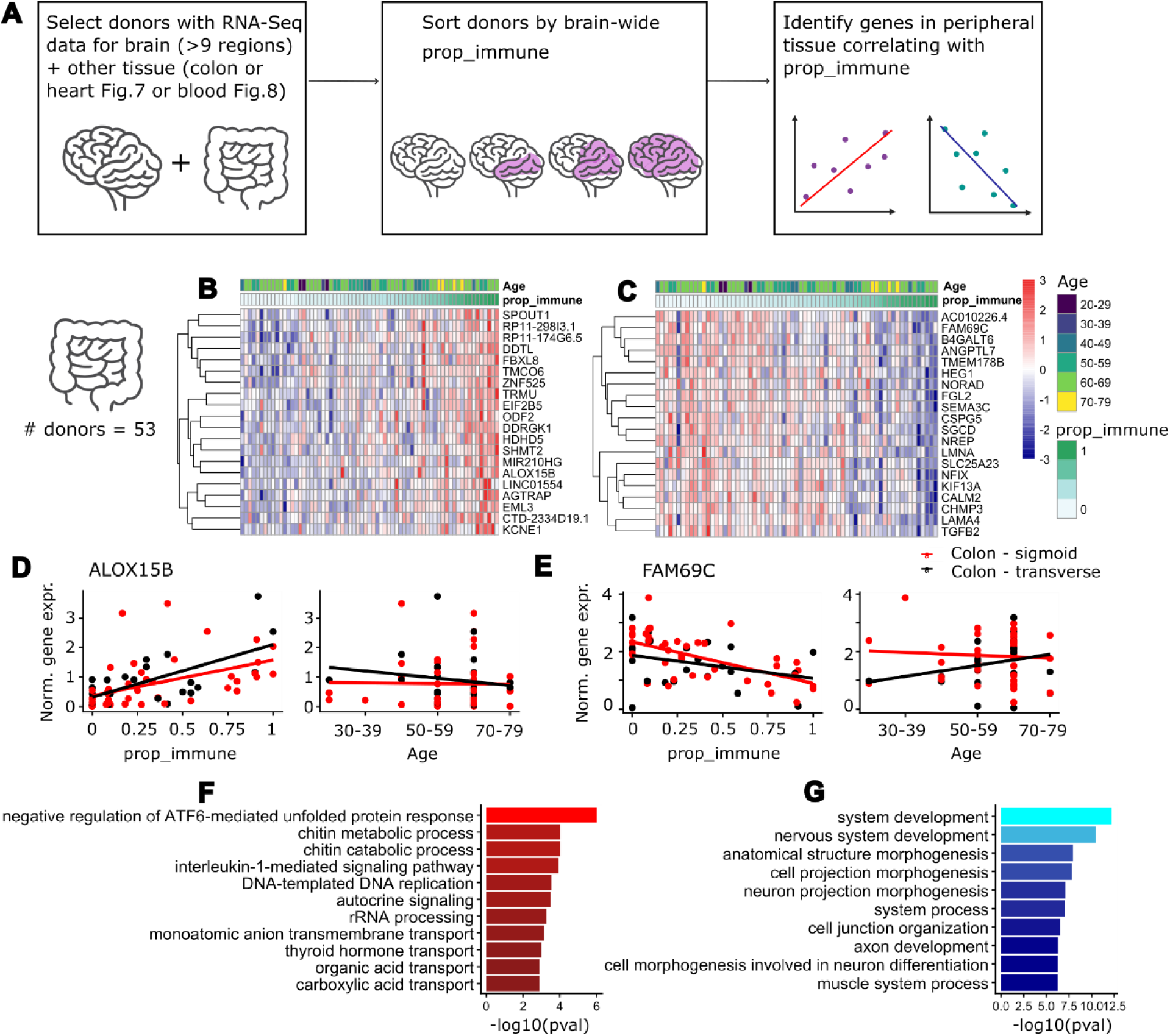
Brain-wide expansion of the immune phenotype is associated with transcriptomic changes in colon **(A)** Methodological workflow illustrating the framework linking the proportion of immune-type samples (prop_immune) to gene expression in peripheral organs within the same donors. **(B and C)** Hierarchical clustering heatmaps of the top 20 transcripts in colonic biopsies demonstrating direct (B) or inverse (C) correlations with increasing brain prop_immune values (n = 53 donors). Matrix values represent scaled, log2-normalized gene expression levels adjusted for age, sex and anatomical tissue subregion (sigmoid vs. transverse colon). **(D and E)** Selected scatter plots showing colon expression values for ALOX15B (D, an increasing gene with a positive slope of 1.8 in colon-sigmoid and 1.2 in colon-transverse) and FAM69C (E, a decreasing gene with a negative slope of -1.3 in colon-sigmoid and -0.77 in colon-transverse) plotted against brain prop_immune (left panels) and donor age (right panels), revealing specificity for prop_immune over general chronological aging. Sigmoid (red) and transverse (black) subregions are tracked independently, highlighting uniform directional slopes across the colon. **(F and G)** GO BP enrichment analysis of colonic transcripts with significant direct (F) or inverse (G) correlation with prop_immune (FDR < 0.05, |Pearson’s r| > 0.3). Bar values denote -log10(p-value).

Pathway enrichment analysis revealed significant upregulation of metabolic and immune-responsive processes, particularly carboxylic acid transport and chitin metabolic pathways, indicative of a shift in colonic epithelial function (Fig. 6F). In parallel, a robust down-regulation of pathways involved in structural integrity was observed, including cell junction organization, muscle system processes, and nervous system development. These alterations point to substantial impairment of neuromuscular and epithelial integrity within the colon (Fig. 6G).

Application of the same analysis to heart tissues (79 donors, atrial appendage n = 67 samples, left ventricle n = 51 samples) revealed a more limited transcriptomic signature with 111 genes significantly correlating with prop_immune and 34 showing inverse correlation (Fig. 7, A and B, Table S1). *S100A8,* a well-established inflammatory biomarker, showed strong correlation with prop_immune and the steepest positive slope (slope = 2.3), whereas *NREP*, a regulator of cardiac tissue remodeling, showed one of most negative values (slope = -1.5) and a significant inverse correlation (Fig. 7, C and D). Pathway enrichment analysis revealed increased inflammatory signaling alongside reduced angiogenic capacity in association with brain-wide expansion of the immune phenotype (Fig. 7, E and F).

**Fig. 7.**
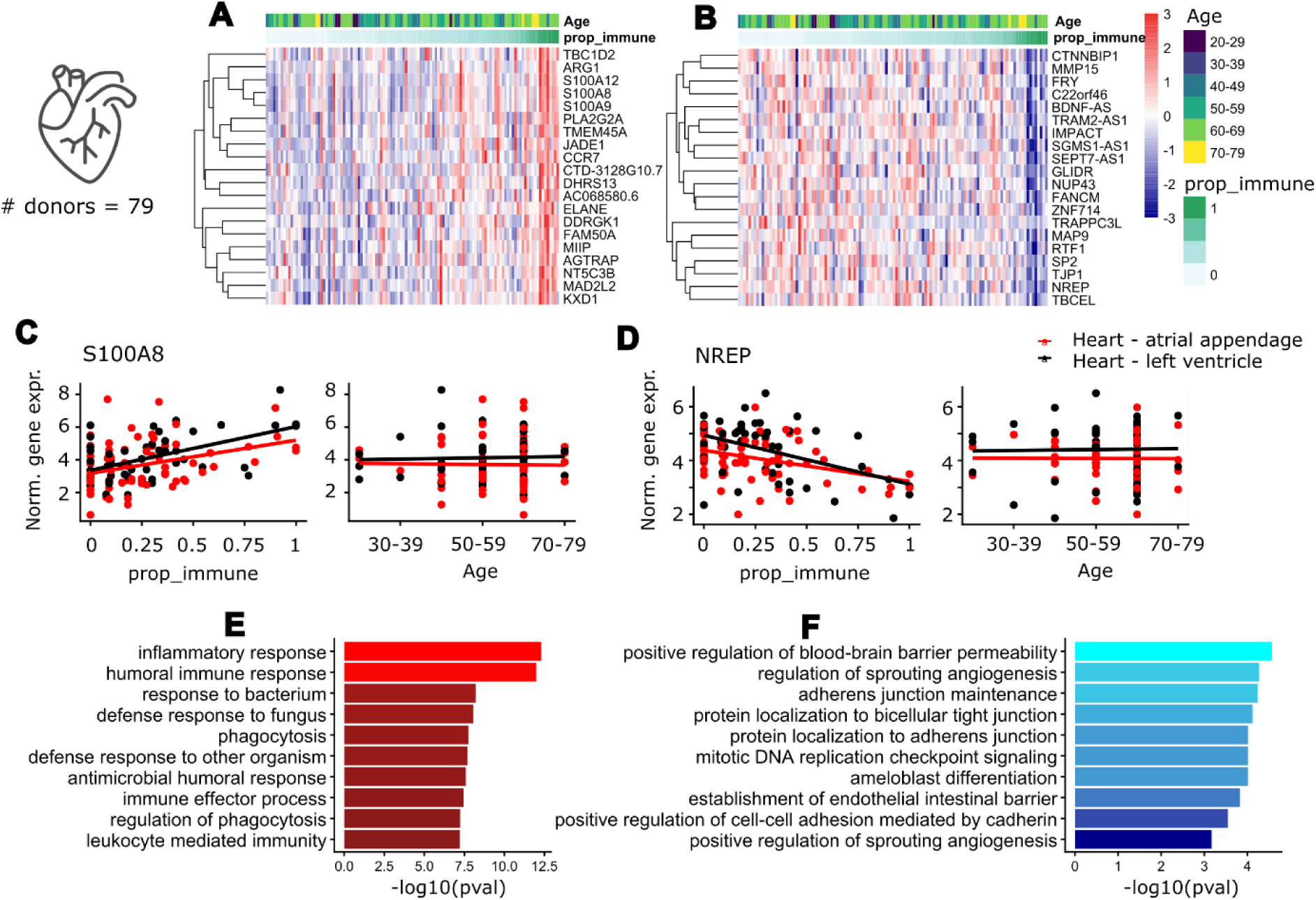
Brain-wide expansion of the immune type is associated with transcriptomic changes in the heart **(A-B)** Hierarchical clustering heatmaps of the top 20 transcripts in cardiac tissues demonstrating direct (A) or inverse (B) correlations with increasing brain prop_immune values (n = 79 donors). Matrix values represent scaled, log2-normalized gene expression levels adjusted for age, sex and anatomical tissue subregion (atrial appendage vs. left ventricle). **(C and D)** Selected scatter plots showing heart expression values for S100A8 (C, an increasing gene with a positive slope of 2.0 in heart-atrial appendage and 2.7 in heart-left ventricle) and NREP (D, a decreasing gene with a negative slope of -1.2 in heart-atrial appendage and -1.8 in heart-left ventricle) plotted against brain prop_immune (left panels) and donor age (right panels). Atrial appendage (red) and left ventricle (black) subregions are tracked independently, highlighting uniform directional slopes across the heart. **(E- F)** GO BP enrichment analysis of cardiac transcripts with significant direct (E) or inverse (F) correlation with prop_immune (FDR < 0.05, |Pearson’s r| > 0.3). Bar values denote -log10(p-value).

Finally, we applied the same analysis to whole blood samples. Transcriptomic data were available for 87 donors for whom a corresponding prop_immune value had been determined. After adjusting for sex and age, we identified 326 genes significantly correlating with prop_immune and 430 genes showing inverse correlation (Fig. 8, A and B, Table S1). *AIM2*, a central regulator of pyroptosis, exhibited strong correlation with prop_immune and one of the steepest positive slopes (slope = 1.1), while *HLA-DQA1*, a central mediator of adaptive immunity, showed one of the lowest values (slope = -1.9) and significant inverse correlation (Fig.8, C and D). Pathway enrichment analysis revealed a marked upregulation of genes involved in chaperone activity, detoxification and pyroptotic inflammation (Fig. 8E). In contrast, pathways negatively associated with prop_immune were enriched for adaptive immune response, with histocompatibility complex class II genes *HLA-DQA1*, *HLA-DQB1*, and *HLA-DMB* among the most strongly down-regulated transcripts (Fig. 8F). These findings suggest that brain-wide expansion of the immune phenotype is associated with increased inflammation accompanied by a reduction in adaptive immune activity in the blood.

**Fig. 8.**
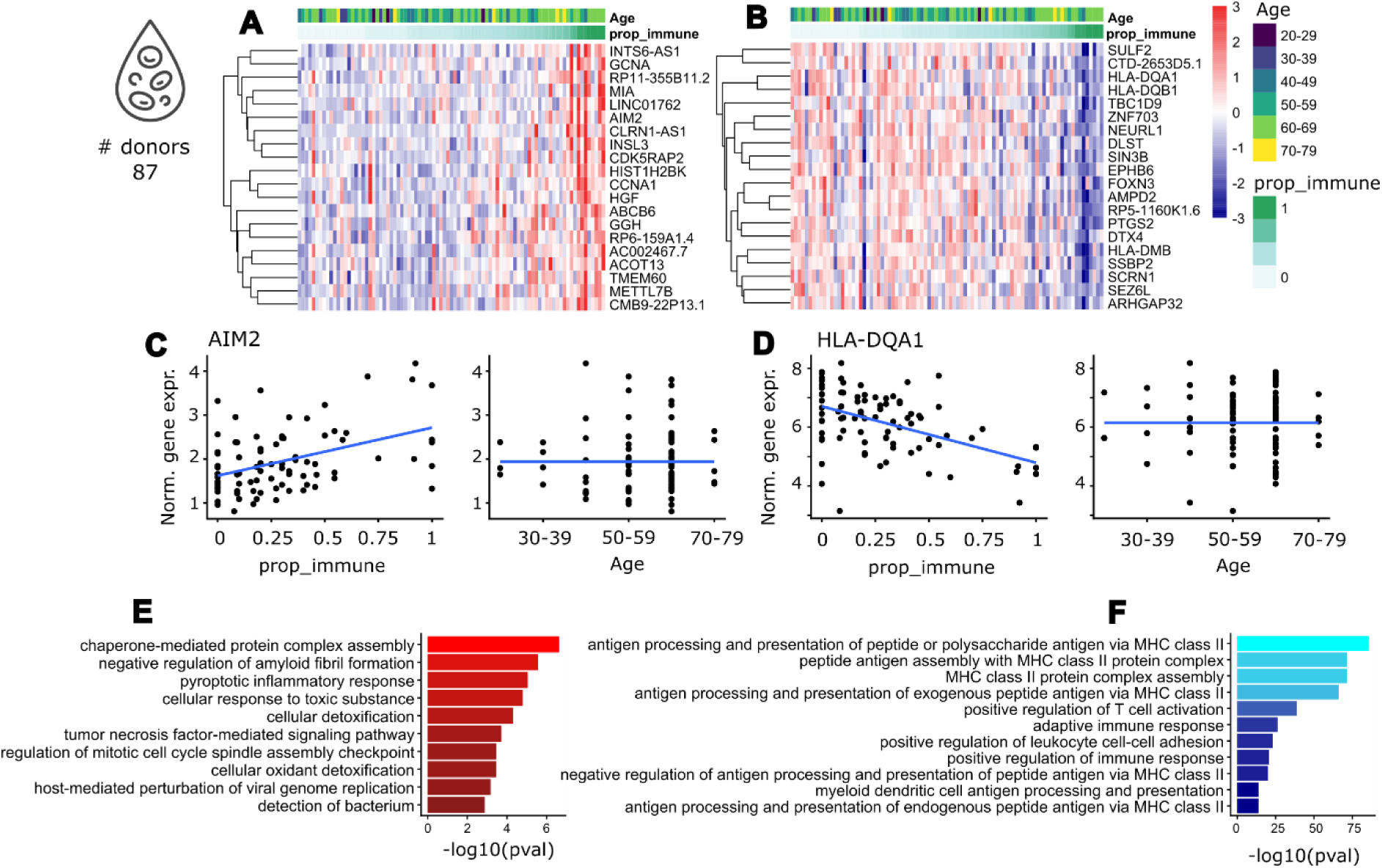
Brain-wide expansion of the immune type is linked to coordinated transcriptomic changes in whole blood **(A and B)** Hierarchical clustering heatmaps of the top 20 whole-blood transcripts demonstrating direct (A) or inverse (B) correlations with increasing brain prop_immune values (n = 87 donors). Matrix values represent scaled, log2-normalized gene expression levels adjusted for age and sex. **(C and D)** Selected scatter plots showing blood expression values for *AIM2* (C, an increasing gene with a positive slope of 1.1) and *HLA-DQA1* (D, a decreasing gene with a negative slope of -1.9) plotted against brain prop_immune (left panels) and donor age (right panels), revealing specificity for prop_immune over general chronological aging. **(E and F)** Gene Ontology Biological Process (GO BP) enrichment analysis of blood transcripts with significant direct (E) or inverse (F) correlation with prop_immune (FDR < 0.05, |Pearson’s r| > 0.3). Bar values denote -log10(p-value).

Together, these results revealed coordinated molecular alterations across multiple peripheral tissues that accompany the topographical expansion of the immune phenotype throughout the CNS. The shared enrichment of inflammatory pathways in the colon, heart, and blood supports the presence of a systemic immune axis linked to the emergence and expansion of the immune phenotype in the brain.

### A conserved systemic inflammatory response associates with brain-wide expansion of the immune phenotype

To identify mechanisms underlying the coordinated immune response across multiple tissues, we defined a shared peripheral and CNS molecular signature associated with the brain-wide expansion of the immune phenotype. To this end, we evaluated the relationship between gene expression and prop_immune across all brain samples, adjusting for sex, age, and brain region, and identified genes significantly correlating with increasing prop_immune (Table S1). Intersecting the results from brain and blood revealed 8 genes strongly correlated with prop_immune in both compartments (FDR < 0.05, Pearson’s r > 0.3, slope > 1): *AGTRAP*, *RAB13*, *BCL6*, *SULT1B1*, *CD163*, *MGST1*, *SERPINA1*, and *ITPKC* (Fig. 9A). These genes are functionally linked to inflammatory signaling pathways and are predominantly expressed in myeloid cell populations, including monocytes, neutrophils, and granulocytes. We next examined the association between the 8-gene signature and prop_immune across all GTEx tissues with at least 20 profiled donors, adjusting for age and sex. Most peripheral tissues demonstrated a consistent positive association between prop_immune and the expression of all 8 genes, supporting the presence of a coordinated systemic inflammatory response associated with the expansion of the immune phenotype in the brain. Finally, we extended this analysis to three additional clinically relevant, inflammation-associated gene sets: canonical inflammatory markers (*CRP* and *CHI3L1*; Set 2); genes modulated by non-steroidal anti-inflammatory drugs (NSAIDs; *PTGS1, PTGS2, IL6, TNF, NFKB1*, and *NFKB2*; Set 3), and inflammasome-related genes (*NLRP3, CASP1, PYCARD, IL1B*, and *IL18*; Set 4) (Fig. 9B). Notably, *CRP*, the prototypical marker of systemic inflammation, was significantly upregulated in peripheral blood and displayed the strongest association with prop_immune in the liver, the primary site of *CRP* synthesis, although this trend did not reach statistical significance. *CHI3L1* expression increased in several tissues but showed inverse correlation with prop_immune in blood. Among NSAIDs responsive genes, *NFKB1, NFKB2, IL6* and *PTGS2* exhibited upregulation across multiple brain areas. The inflammasome-associated gene *CASP1* was also significantly upregulated in several brain regions, while *NLRP3* was significantly downregulated. *IL18* and *PYCARD* showed increased expression only in three brain regions, the hippocampus, the caudate, and the anterior cingulate cortex. *PTGS1, TNF*, and *IL1B* were not significantly associated with prop_immune. These findings suggest engagement of NSAID-modulated inflammatory pathways during brain-wide expansion of the immune phenotype, while providing limited evidence for sustained activation of the canonical inflammasome pathway.

**Fig. 9.**
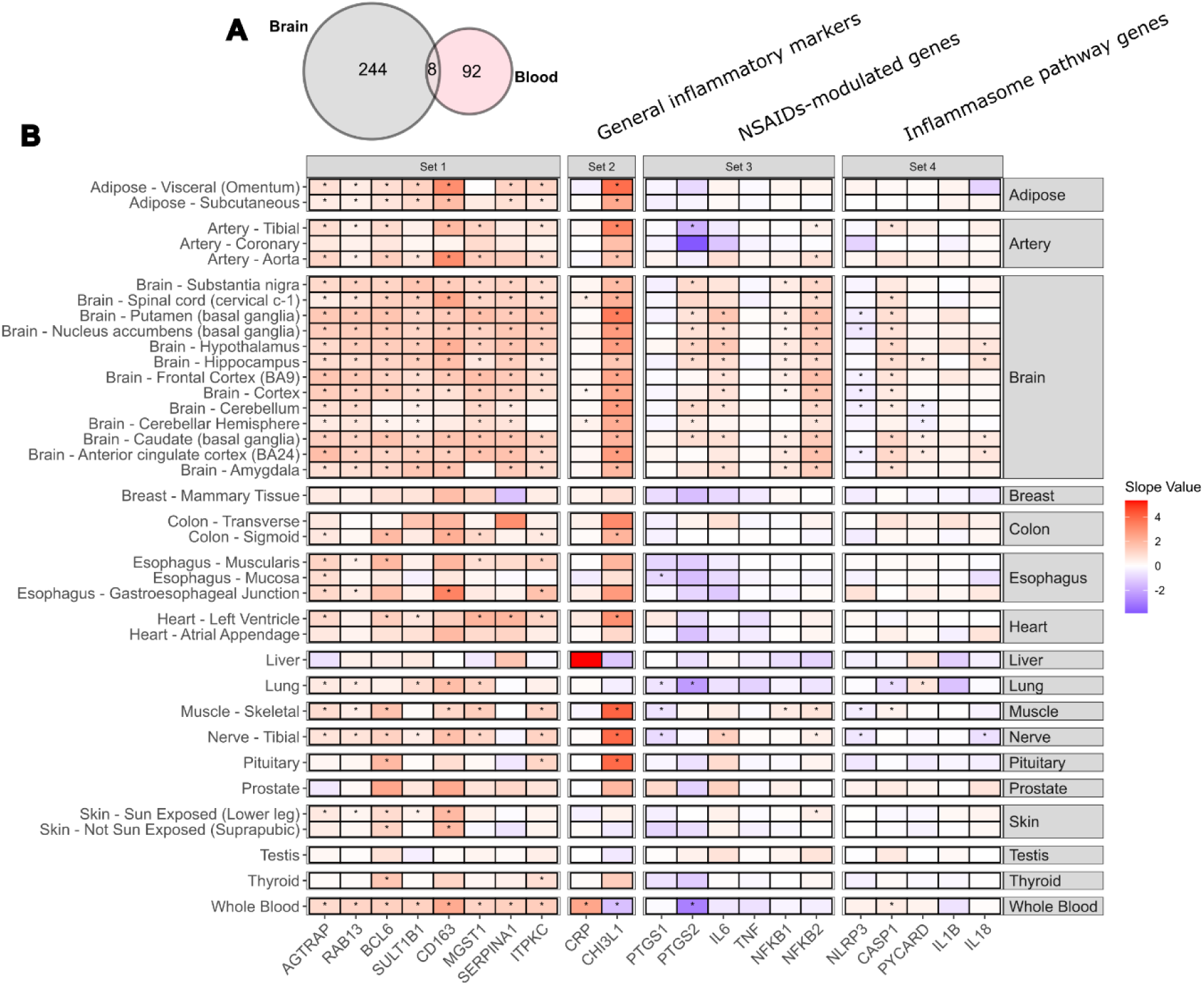
Identification of a shared CNS-peripheral immune signature and its multiorgan expression landscape **(A)** Venn diagram illustrating the intersection of genes significantly correlated with prop_immune in the whole brain (gray) and peripheral blood (pink). A core set of 8 overlapping genes (AGTRAP, RAB13, BCL6, SULT1B1, CD163, MGST1, SERPINA1, and ITPKC; Set 1) was identified based on strict filtering criteria (FDR < 0.05, correlation coefficient > 0.3, slope > 1). **(B)** Heatmap displaying the tissue-specific regression slope values for the 8 signature genes (Set 1), general inflammatory markers CRP and CHI3L1 (Set 2), NSAIDs modulated genes PTGS1, PTGS2, IL6, TNF, NFKB1 and NFKB2 (Set 3), inflammasome pathway genes NLRP3, CASP1, PYCARD, IL1B and IL18 (Set 4) across various central and peripheral organs with data available for at least 20 donors. Gene expression was adjusted for sex and age before calculating the linear model. Color intensity represents the slope value, ranging from downregulation (blue) to upregulation (red). Asterisks indicate Benjamini- Hochberg false discovery rates: *FDR< 0.05.

Taken together, the multi-organ transcriptomic analyses demonstrated that the development of a consistent immune phenotype throughout the brain is accompanied by a systemic inflammatory state spanning peripheral tissues and the CNS. Moreover, the consistent upregulation of NSAIDs-responsive genes across multiple brain regions identifies inflammatory pathways potentially amenable to therapeutic modulation. These findings raise the possibility that early modulation of NSAID-responsive inflammatory pathways may reduce the likelihood of expansion of the immune phenotype in the brain, supporting further investigation of NSAIDs as potential preventive interventions in susceptible individuals.

## DISCUSSION

Several recent studies have focused on dissecting the molecular heterogeneity of patients with neurodegenerative diseases and proposed the existence of distinct subtypes of AD (17, 18), Parkinson’s disease (21), multiple sclerosis (22, 23), amyotrophic lateral sclerosis (24) and other disorders (25). However, these analyses have generally assumed that the identified molecular programs represent disease-specific subtypes. Our results challenge this interpretation, demonstrating that brain subtypes previously described as AD-specific exist also in neurologically healthy individuals. Indeed, in analyzed brain datasets, subtype classification accounted for greater transcriptional variation than AD neuropathology or cognitive decline, indicating that these molecular programs capture an underlying phenotype of the brain rather than consequences of disease. This observation is consistent with recent analysis of cerebrospinal fluid proteome revealing comparable molecular subtypes in healthy individuals and Alzheimer’s patients (26, 27). Furthermore, an APOE ε4-associated immune signature was identified across multiple neurodegenerative diseases and neurologically healthy individuals, with corresponding changes in the brain, CSF and plasma (19). Similarly, assessment of transcriptomic heterogeneity in the brain of patients with multiple sclerosis showed that patient-specific molecular variability exceeded lesion-associated transcriptional changes (22). These findings suggest that at least some molecular subtypes described across neurodegenerative diseases may instead represent brain phenotypes that are autonomous yet interacting with disease, potentially modulating its onset or progression. This distinction has important implications. In our study, approximately 70% of genes identified as differentially expressed in AD overlapped with the signature of the immune phenotype. Accordingly, stratification of samples by immune-type classification markedly reduced the number of AD-associated transcriptional changes, indicating that a substantial fraction of the AD signature might be influenced by the underlying immune phenotype. Similar observations have recently been reported in schizophrenia, where most disease-associated mRNA differences disappeared after stratification of samples according to the inflammatory profile (28). Therefore, comparisons between molecularly heterogeneous disease and control cohorts may primarily capture differences in brain phenotypes rather than disease-specific mechanisms. We propose that future studies should monitor donor-to-donor heterogeneity and, when necessary, stratify both cases and controls according to the underlying molecular brain phenotype before defining disease signatures or prioritizing therapeutic targets.

Several factors likely contribute to shaping long-term brain phenotypes, including cardiovascular disease (29, 30), metabolic dysfunction (31, 32), obesity (33, 34), chronic infections (35), environmental exposures (36, 37) and genetic background (38, 39). Limited available metadata prevented us from performing a thorough investigation of the different factors and clinical outcomes associated with the immune phenotype. Instead, we could leverage a rich collection of transcriptomic data to demonstrate a connection between the immune brain phenotype, age and peripheral inflammation. A major novelty of this study is the use of several brain areas to assess inter-donor heterogeneity, encompassing cortical regions, basal ganglia, limbic areas, midbrain and cerebellum. Analyzing multiple brain areas from the same neurologically healthy donors, we discovered that the cross-regional consistency of the immune type varies substantially among individuals and gradually increases with age. This suggests that rather than representing a fixed property established early in life, a brain-wide consistent immune phenotype might develop over time. We therefore proposed that the proportion of brain regions exhibiting the immune phenotype – a parameter we called prop_immune – could represent the potential “stage” of progression towards a brain-wide consistent immune phenotype. Analyzing transcriptional changes associated with prop_immune in the brain and peripheral organs, we identified coordinated variations in gene expression in the CNS and periphery, with a conserved inflammatory gene signature shared across organs. These observations argue against the immune phenotype representing an isolated neuroinflammatory process and support the pivotal role of systemic inflammation in its development. The pattern of spatial expansion of the immune phenotype within the brain further supports the possibility that peripheral processes contribute to its establishment. Although our cross-sectional analyses cannot establish temporal propagation, the observed topography is compatible with a model in which a peripheral trigger progressively engages the spinal cord and brainstem before extending along mesolimbic and mesocortical pathways towards the cortex. Such a model is reminiscent of the body-first hypothesis proposed for Parkinson’s disease (40, 41), where the pathology originates in the gut. Our analysis confirms that individuals with higher prop_immune present alterations in genes involved in maintaining the structural integrity of the enteric system in the colon. This is consistent with growing evidence linking gut dysbiosis, microbial products and enteric pathogens to blood-brain barrier dysfunction, neuroinflammation, and neurodegeneration (42).

Interestingly, the topographical expansion of the immune phenotype within the brain, was not associated with the induction of canonical inflammatory mediators such as *IL1B* or *TNF,* or inflammasome. Instead, increased expression of *PTGS2*, *IL6*, *NFKB1*, and *NFKB2* suggests that brain-wide expansion of the immune phenotype might be driven by a selective chronic inflammatory program, which could be targeted by non-steroidal anti-inflammatory drugs (NSAIDs). Curiously, the proportion of neurologically healthy donors with a highly consistent immune-type classification across brain areas (6 – 22% with prop_immune ≥ 0.75, depending on age) closely resembles the reduction in AD incidence (12 – 20%) reported in epidemiological studies of NSAIDs use (43–45). Although speculative, this correspondence raises the possibility that the cohort that showed decreased AD risk following NSAIDs treatment in previous prevention studies was composed of individuals with the immune brain phenotype. Prospective studies combining transcriptomic biomarkers with preventive intervention will be required to test this hypothesis.

Although the immune phenotype exists independently of AD, several observations indicate that it influences disease susceptibility. First, immune-type samples were significantly more frequent among AD donors and APOE ε4 carriers. Second, immune-type samples were enriched in reactive microglia, including a microglia subpopulation previously associated with Aβ pathology (CPM^+^ Mic.12) and were depleted in neurons, specifically SST^+^ interneurons which are the most vulnerable in AD (20). Third, immune-type PCC or TC was preferentially associated with cognitive impairment, whereas an immune-type DLPFC or FC was associated with neuropathological burden. Interestingly, these regional differences mirror the functional organization of large-scale brain networks and suggest that the biological consequences of the immune phenotype depend on the circuits in which it develops. Both PCC and TC serve as critical core hubs of the Default Mode Network (DMN), a system directly tied to episodic memory collapse (46). Conversely, the DLPFC is a primary component of the Central Executive Network (CEN), which is characteristically affected at much later stages of AD tissue pathology (47). Therefore, the occurrence of the immune phenotype within the CEN indicates greater AD neuropathology burden across the brain, but the occurrence of the immune phenotype in the DMN enhances the risk of overt cognitive failure. Most notably, immune-type DLPFC was specifically associated with increased Aβ deposition but not tau pathology. While control donors with little or no AD neuropathology displayed an age-dependent increase in neocortical Aβ levels when exhibiting an immune-type DLPFC, this association was absent in AD cases. These characteristics align with the antimicrobial protection hypothesis (48), which describes Aβ aggregation as a form of protection of the brain against microbes and therefore proposes that infection and inflammation could be the initial triggers of the amyloid cascade. Future investigations should explore whether exposure to specific viruses and pathogens could drive the processes leading to the development of the immune brain phenotype. Despite its association with Aβ accumulation, the immune-type DLPFC was not associated with altered cognitive decline at any stage of neuropathological progression. Together, these observations suggest that the immune phenotype primarily modulates disease initiation or susceptibility rather than severity of disease progression. Longitudinal studies monitoring biomarkers of the immune phenotype with blood levels of pTau-217 might provide evidence supporting this hypothesis.

In summary, our study supports a revised framework for understanding molecular heterogeneity in neurological disorders. Our findings indicate that the immune-type classification captures a distinct brain phenotype that develops independently of AD, is associated with increased cortical Aβ accumulation, and coincides with coordinated inflammatory changes across peripheral tissues.

### Limitations

The first limitation of our study is the exclusive usage of post-mortem samples, allowing us to perform only cross-sectional analysis. As pointed out in several parts of the discussion, longitudinal measurements are required to confirm age-dependent acquisition of a brain-wide consistent immune phenotype and its association with progressive Aβ accumulation. Longitudinal analysis of blood samples could also facilitate biomarker discovery to reliably identify immune-type individuals in the clinic. The second major limitation is our limited access to metadata and whole genome sequencing data for most donors analyzed. A follow-up study will consider partnership with consortia to access protected data and investigate genetic risk and comorbidities associated with the immune phenotype.

## Material and Methods

### Cohorts and Data Sources

The current study utilizes data from 4 sources: the Genotype-Tissue Expression (GTEx) project, the Religious Orders Study and Memory and Aging Project study (ROSMAP), the Mount Sinai Brain Bank (MSBB) study and the MAYO RNA-Seq Study (MAYO). Besides GTEx, all other studies are part of the AMP-AD consortium. No new data are generated but secondary analysis of the de-identified patient data is performed. Here we provide a brief description of the cohorts and data types used for analysis in this study, while detailed metadata and graphical summary of cohorts composition are provided in Tables S2 and Fig. S2, respectively.

GTEx: A comprehensive resource collecting data from up to 54 non-diseased tissue sites across ∼1000 deceased individuals, covering an age range of 20 to 79 years of age. Bulk transcriptomics data have been generated for several tissues and donors, allowing a comprehensive analysis at the individual levels of molecular signatures throughout the body. In this study, we use a subset of GTEx bulk transcriptomics resource: all CNS (brain + spinal cord), heart and colon samples published within version 8 (49). Demographic and clinical data are available online and can be freely downloaded from the GTEx website.

ROSMAP: Consists of two longitudinal, clinicopathological, community-based cohort studies, ROS and MAP, initiated in 1994 and 1997 respectively. Both studies recruit participants without known dementia in the United States and involve annual clinical evaluations of cognitive performance and post-mortem brain donation. The Rush University Medical Center’s Institutional Review Board approved the ROSMAP study and all participants provided informed consent. Bulk transcriptomics data have been generated for a total of 1148 donors, from samples of dorsolateral prefrontal cortex (DLPFC, n = 1091), posterior cingulate cortex (PCC, n = 657), head of caudate nucleus (HCN, n = 726), temporal cortex (TC, n = 125) and frontal cortex (FC, n = 123). Single nuclei transcriptomics data were generated from DLPFC samples for a total of 465 donors, reduced to 437 after QC and elimination of samples with insufficient number of nuclei, as described in syn31512863. The list of nuclei belonging to the 437 participants and used in the current study for abundance analysis is available under cell-annotations.n437.csv (<u>syn53694215</u>). Demographic and clinical data stored in <u>syn27000096</u> are used in this study and explanation of each variable is provided at the www.radc.rush.edu website. Quantification of Aβ and tangles and calculation of the rate of cognitive decline in a subset of donors was performed in a separate study (20) and can be found in the corresponding supplementary materials. Briefly, the rate of cognitive decline was calculated by longitudinally modelling the annual global cognitive scores with a mixed effects model, adjusting for age, sex and education, providing a slope of decline for each participant. The burden of parenchymal Aβ and tangles of hyperphosphorylated tau was quantified by immunohistochemistry in 8 brain regions (hippocampus, entorhinal cortex, anterior cingulate cortex, midfrontal cortex, superior frontal cortex, inferior temporal cortex, angular gyrus and calcarine cortex) and summarized to provide a global measure of Aβ and tau burden.

MSBB: Comprises several human brains and different brain regions, obtained from the Mount Sinai/JJ Peters VA Medical Center Brain Bank and provides multi-omics data across a spectrum of AD severity, ranging from severe cognitive impairment and control cases. Bulk transcriptomics data have bene generated for a total of 315 donors, from samples of parahippocampal gyrus (PHG, n = 310), superior temporal cortex (STC, n = 327), inferior frontal cortex (IFC, n = 301) and frontal pole (FP, n = 306). This study utilizes a subset of the entire collection for two specific analyses: 214 PHG samples are used to replicate previously published analysis (18). 272 STC samples (one sample per donor in the dataset, excluding replicates) are used to further corroborate the independence of clusters to AD diagnosis in a separate collection of data. Demographic and clinical data stored in <u>syn6101474</u> are used in this study. This version (version 10) of clinical data has been reviewed and corrected of the discrepant entries identified in August 2025 among different versions of the metadata.

MAYO: Comprises several human brains and different brain regions, obtained from the Mayo Clinical Brain Bank (MCBB) or the Banner Sun Health Research Institute (Banner). Bulk transcriptomics data have been generated for a total of 312 participants with AD, progressive supranuclear palsy (PSP), pathologic aging (PA) or elderly controls (CTR) without neurodegenerative diseases, from samples of temporal cortex (TC, n = 276) and cerebellum (CBE, n = 275). This study utilizes only TC samples from AD, PA and CTR donors and excludes PSP from the analysis. Demographic and clinical data stored in <u>syn26403544</u> are used in this study.

### Harmonization of Metadata in AD Cohorts

The diagnosis criteria for AD differ across ROSMAP, MSBB and MAYO datasets. To harmonize the classification of patients, we defined three types of diagnosis and applied the same criteria to all datasets. When not enough information was available to attribute a clear diagnosis based on the selected criteria, the diagnosis was annotated as “unknown”. The three diagnosis fields are the following. 1) Clinical diagnosis: refers to the clinical symptoms, reported in MSBB with CDR score and in ROSMAP with cogdx score. We assign the following levels: CTR (CDR =0 or cogdx = 1), MCI (CDR = 0.5 or 1, cogdx = 2 or 3), AD (CDR = 2 or 3, cogdx = 4 or 5), Very Severe AD (CDR = 4 or 5), Other Dementia (cogdx = 6). MAYO reports directly a dichotomized clinical diagnosis of CTR or AD. 2) Neuropathological diagnosis: based on the topographic expansion of Aβ and Nft in the brain, uses an approximation of the ABC score (50) to assign a diagnosis of AD. Most donors lacked Thal phase, so we assigned the following levels: CTR (B_score = 0 or 1; B_score = 2 and C_score = 0, MAYO CTR donors), AD (B_score = 2 or 3 and C_score = 2 or 3, all MAYO AD and PA donors), unknown (B_score = 2 or 3 and C_score = 1). 3) Joint diagnosis: refers to the combination of clinical and neuropathological diagnosis and has the following levels (clinical_neuropathological): CTR (CTR_CTR), MCI- (MCI_CTR), MCI+ (MCI_AD), AD (VerySevereAD_AD or AD_AD), resilient (CTR_AD), Other Dementia (Other Dementia_CTR or AD_CTR or VerySevereAD_CTR), unknown (Other Dementia_AD or missing values in one diagnosis field).

### Weighted Gene Co-expression Network Analysis (WGCNA)

The R package WGCNA (51, 52) was used to construct a co-expression network using the bulk transcriptomics data of ROSMAP, MSBB and MAYO, temporal cortex samples. Count matrix was first normalized to Counts Per Million (CPM) values using Trimmed Mean of M-values (TMM) method, log2 transformed and adjusted for sex, study and batch using limma R package. The weighted adjacency matrix was constructed by calculating the pairwise Pearson correlation coefficients between all genes and applying a soft-thresholding power of 6 (as the smallest threshold resulting in a scale-free fit index R^2^ of ∼0.9 and mean connectivity ∼90). Modules were identified using hierarchical clustering based on the Topological Overlap Measure (TOM) followed by a dynamic tree-cut algorithm (minimum module size = 30). 20 modules were identified and numbered according to size, starting with the largest (M1, n = 6950). To determine the association between diagnosis and each module, the first principal component of the module’s gene expression, called eigengene, was used.

### Identification and Definition of Clusters

To identify clusters of samples we used the WSCNA algorithm (18), which treats samples as nodes in a weighted network to evaluate their global transcriptomic similarity. The algorithm is based on a computationally optimized version of WGCNA (53), using the same mathematical framework, but transposes the input matrix (columns represent genes and rows samples) so that sample-sample correlations are compared. Count matrix was first normalized to Counts Per Million (CPM) values using Trimmed Mean of M-values (TMM) method, log2 transformed and adjusted for covariates (specified for each analysis in the results section) using limma R package. Parameters used: soft-thresholding power = 5.9, minimum module size = 50.

The signature of each cluster was defined as the genes differentially expressed in one cluster compared to all others pooled together.

### Differential Gene Expression Analysis

Differential gene expression (DEG) analysis was performed in R using the package limma. The count matrix was first normalized to CPM values using TMM method, lowly expressed genes were excluded (max CPM < 3) and finally values were transformed using voom (limma R package). Gene expression values for samples in the comparisons were then fit to a linear model (lmfit, limma R package), using a design that removes the effect of confounding variables. Confounding variables depend on the dataset and the analysis performed and are specified in the results section. Contrasts were fit and empirical Bayes statistics for differential expression were calculated (eBayes, R function), followed by the topTable R function to output significant DEGs. Adjusted p-values were calculated using the standard Benjamini-Hochberg (BH) correction for multiple comparisons. Signatures were defined by genes with adjusted p-value < 0.05 and absolute value of logFC > 0.1.

### Comparison of Clusters

Clusters’ signatures were compared using a modified Jaccard index. The modified Jaccard Index quantifies the similarity between a pair of clusters, considering the coherence in the direction of regulation of each gene:

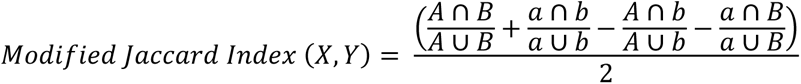

For each pair of clusters (X,Y), we define A and B as up-regulated genes and a and b as down-regulated genes in the first (X) and second (Y) cluster respectively.

### Pathway Enrichment

Gene Set Enrichment Analysis was performed using the evoGO package in R (54), developed by Evotec International GmbH. The evoGO algorithm addresses the issue of redundancy in classic GO overrepresentation analysis and reduces the significance of less specific terms if their DEGs contribute to higher significance in descendent terms. EvoGO was applied on up-regulated and down-regulated DEGs separately to define up and down-regulated terms respectively. An evoGO p-value < 0.05 was used to define significant GO terms.

### Definition of Cell Types Proportions in ROSMAP snRNA-Seq data

Annotation of cell types and cell type subpopulations was taken from Green et al., work on ROSMAP DLPFC snRNA-Seq data (20). Our analysis focuses on 11 major cell type groups (excitatory neurons, inhibitory neurons, astrocytes, microglia, monocytes, macrophages, oligodendrocytes, OPCs, Endothelial, Pericytes and SMCs) and the subpopulations of neurons, microglia and astrocytes. We decided not to focus on subpopulations of oligodendrocytes, OPCs and vascular niche cell types because of the difficulty in interpretation of the different subpopulations/ states associated to these cell types. For each sample, the proportion of each major cell type was calculated by dividing the number of nuclei belonging to each cell type by the total number of nuclei in the sample. The subpopulations proportions were then calculated within each relevant cell type group by dividing the number of nuclei of each subpopulation by the total number of nuclei of the corresponding cell type group.

### Inferring Cell Types Proportion from Bulk RNA-Seq Data

To infer cell types and subpopulations proportions, we used the CelMod method [https://github.com/MenonLab/Celmod (20, 55)]. Compared to other deconvolution methods that require a pre-defined gene signature for each cell type, CelMod has the advantage to calculate the relationship between cell type proportions and gene expression, from a training set of matching bulk and snRNA-Seq data. As training set, we used 75% of ROSMAP DLPFC samples with matching bulk and snRNA-Seq data, while the remaining 25% were used to assess the reliability of the predictions. To reduce sample bias, the algorithm was run 10 times on a random selection of train and test sets, resulting in 10 models and multiple predictions for several samples. We then averaged the predicted cell types and subpopulations proportions, obtaining a single estimated proportion for each sample. The averaged predicted cell types and subpopulations proportion was then compared to the corresponding snRNA-seq proportion in each sample using a Spearman correlation. Only cell types and subpopulations for which correlation was positive and FDR < 0.01 were considered to be reliable and used in downstream analysis. The 10 models trained on matching bulk and snRNA-Seq data were then applied to the remaining ROSMAP DLPFC samples with only bulk RNA-Seq data as well as to GTEx DLPFC (FCx – BA9) bulk RNA-Seq data. For each sample with only bulk RNA-Seq data, 10 predictions of cell types and subpopulations proportions (one for each model) were obtained. The average of the 10 outputs was used in downstream analysis.

To mitigate potential technical biases deriving from the application to GTEx data of a model trained on ROSMAP data, we applied the following validation steps. First, we processed the count matrices of both datasets using the same pipeline: the count matrix is first normalized to CPM values using TMM method, lowly expressed genes are excluded (max CPM < 3); then we confirmed consistency of gene expression between datasets (Fig. S3, A-D). Second, after CelMod application, we verified that the inferred proportion of major cell types is consistent between datasets, with some changes in relative abundance of neurons and glia expected in ROSMAP due to the older age of donors compared to GTEx and the inclusion of several patients with AD (Figure S3, E-G). Third, we compared changes in cellular composition in GTEx CelMod estimates, ROSMAP CelMod estimates, and ROSMAP snRNA-seq– derived proportions, assessing reproducibility of cellular composition of the immune type across datasets and modalities (Figure S4, B-F).

### Statistical Analysis of Compositional Changes in Cell Types Proportions

To compare proportions of cell types or proportion of donors between groups of samples we used a quasi-binomial regression analysis (glm, R function) with confounding variables included in the design and specified in the text. Adjusted p-values were calculated using the standard Benjamini-Hochberg (BH) correction for multiple comparisons.

To assess changes in proportions of cell types across a continuous variable (prop_immune), we used a linear model (lm, R function) and a partial Pearson’s correlation test (pcor_test, R function), adjusting for sex and age. Adjusted p-values were calculated using the standard Benjamini-Hochberg (BH) correction for multiple comparisons.

### Correlation of gene expression with prop_immune

To define changes in gene expression across prop_immune, we initially adjusted gene expression for covariates using limma R package. We then calculated Pearson’s correlation coefficients and p-values (cor.test, R function) and the slope of a linear model (lm, R function) for each gene. Adjusted p-values were calculated using the standard Benjamini-Hochberg (BH) correction for multiple comparisons.

## Acknowledgments

Author contributions

Conceptualization: GMS Methodology: GMS, DR, GGDB, MS Investigation: GMS

Formal analysis: GMS, DR

Data curation: GMS, DR, GGDB Software: DR, GGDB Validation: DR, GGDB, MS

Visualization: GMS, DR, GGDB, MS Resources: GMS, DR, GGDB Funding acquisition: DGS

Project administration: GMS, DGS Supervision: DGS Writing—original draft: GMS, DGS

Writing—review and editing: All authors

## Data Availability

The results published here are based on data obtained from the AD Knowledge Portal (https://adknowledgeportal.org) and The Genotype-Tissue Expression (GTEx) Project. For replication of MSBB PHG clustering, we used the covariates adjusted count matrix stored in <u>syn16795937</u>. Bulk RNA-Seq data of ROSMAP, MSBB and MAYO cohorts are part of the **<u>The</u> <u>RNAseq Harmonization Study (rnaSeqReprocessing)</u>**. Count matrices stored in <u>syn26720675</u> are utilized in this study. Data generation was supported by the following NIH grants: P30AG10161, P30AG72975, R01AG15819, R01AG17917, R01AG036836, U01AG46152, U01AG61356, U01AG046139, P50 AG016574, R01 AG032990, U01AG046139, R01AG018023, U01AG006576, U01AG006786, R01AG025711, R01AG017216, R01AG003949, R01NS080820, U24NS072026, P30AG19610, U01AG046170, RF1AG057440, and U24AG061340, and the Cure PSP, Mayo and Michael J Fox foundations, Arizona Department of Health Services and the Arizona Biomedical Research Commission. We thank the participants of the Religious Order Study and Memory and Aging projects for the generous donation, the Sun Health Research Institute Brain and Body Donation Program, the Mayo Clinic Brain Bank, and the Mount Sinai/JJ Peters VA Medical Center NIH Brain and Tissue Repository. Data and analysis contributing investigators include Nilüfer Ertekin-Taner, Steven Younkin (Mayo Clinic, Jacksonville, FL), Todd Golde (University of Florida), Nathan Price (Institute for Systems Biology), David Bennett, Christopher Gaiteri (Rush University), Philip De Jager (Columbia University), Bin Zhang, Eric Schadt, Michelle Ehrlich, Vahram Haroutunian, Sam Gandy (Icahn School of Medicine at Mount Sinai), Koichi Iijima (National Center for Geriatrics and Gerontology, Japan), Scott Noggle (New York Stem Cell Foundation), Lara Mangravite (Sage Bionetworks). ROSMAP single nuclei annotations part of the **<u>The Religious Orders Study and</u> <u>Memory and Aging Project (ROSMAP) Study</u>** and stored in <u>syn53694215</u> are utilized. Data were generated from postmortem brain tissue provided by the Religious Orders Study and Rush Memory and Aging Project (ROSMAP) cohort at Rush Alzheimer’s Disease Center, Rush University Medical Center, Chicago. This work was funded by NIH grants U01AG061356 (De Jager/Bennett), RF1AG057473 (De Jager/Bennett), and U01AG046152 (De Jager/Bennett) as part of the AMP-AD consortium, as well as NIH grants R01AG066831 (Menon) and U01AG072572 (De Jager/St George-Hyslop). <u>GTEx</u> Bulk RNA-Seq data were obtained from the open source version 8 at the GTEx Portal. The Genotype-Tissue Expression (GTEx) Project was supported by the <u>Common Fund</u> of the Office of the Director of the National Institutes of Health, and by NCI, NHGRI, NHLBI, NIDA, NIMH, and NINDS.

---

**Fig. S1.**
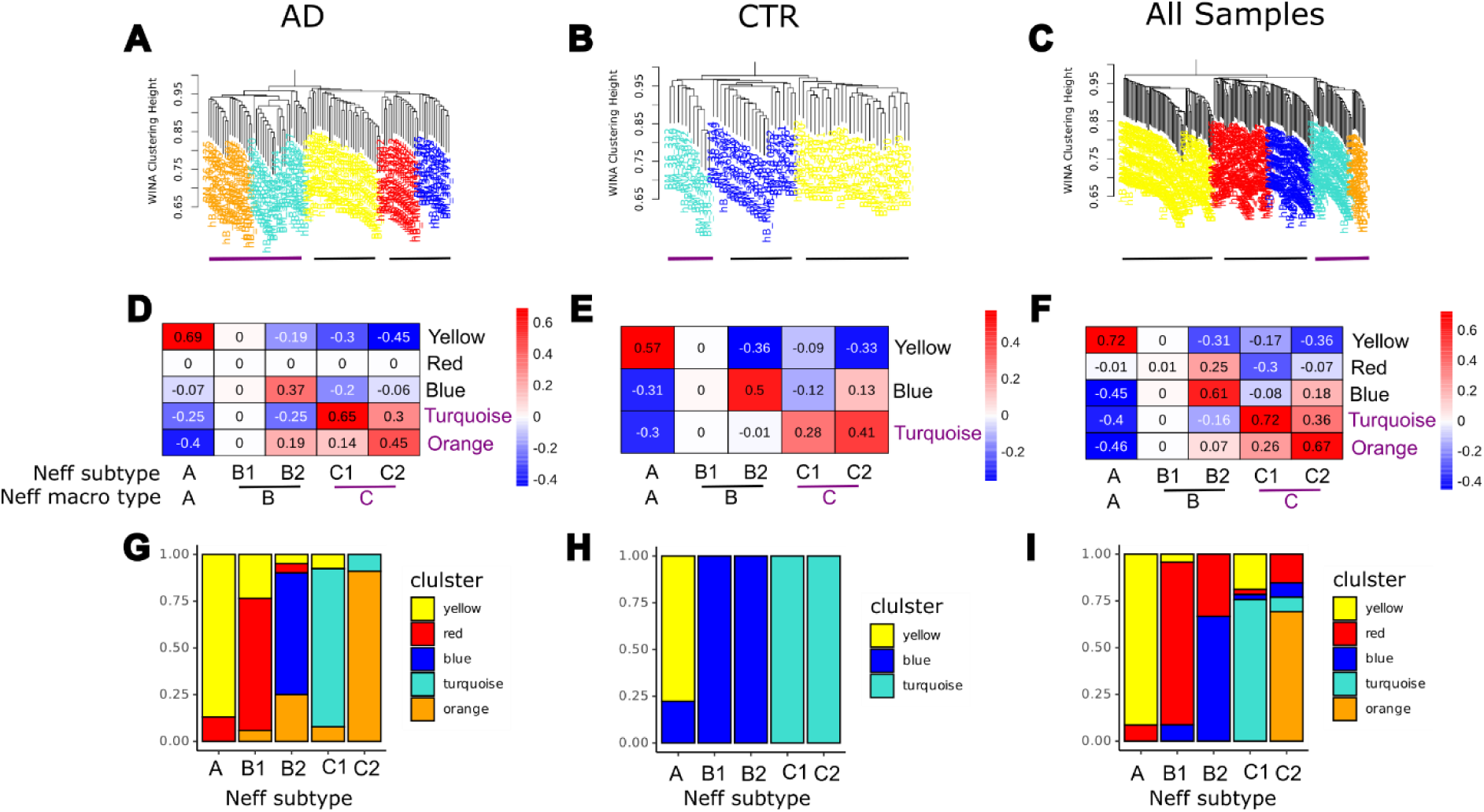
Validation and robustness of WSCNA clustering and module annotation across stratified patient cohorts **(A to C)** Weighted Sample Gene Network Analysis (WSCNA) clustering dendrograms generated from the MSBB parahippocampal gyrus (PHG) dataset, stratified by donor neuropathological diagnosis into three independent groups: AD (A), CTR (B) and all profiled samples combined (C). Horizontal lines at the baseline of each dendrogram delineate the resolved macro-clusters. **(D to F)** Heatmaps depicting the pairwise Jaccard similarity index of characteristic gene signatures between newly resolved sample clusters (rows) and reference subtypes established by Neff et al., 2021 (18) (columns). Matrix values and color bars report the strength of transcriptomic concordance across the AD (D), CTR (E), and combined (F) cohorts. **(G to I)** Stacked bar charts assessing the structural classification consistency between historical annotations and newly derived WSCNA modules across the corresponding cohorts. Analysis is strictly restricted to samples possessing a definitive module assignment in the original reference study.

**Fig. S2.**
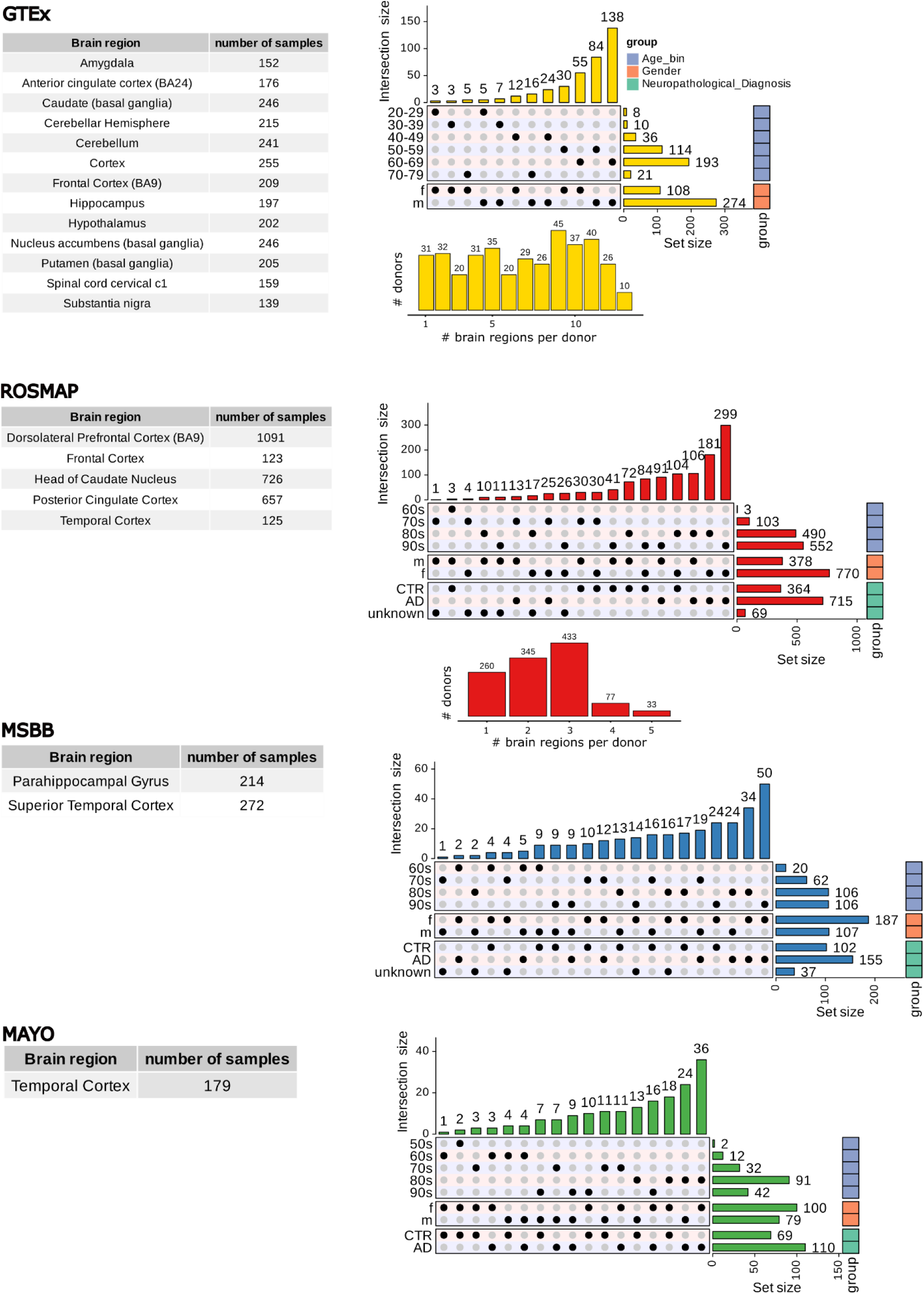
Clinical-pathological metadata and sample distributions across independent brain cohorts Overview of sample sizes, donor counts, and clinical-demographic metadata stratifications across the GTEx, ROSMAP, MSBB, and MAYO cohorts. Left panels detail the exact sample counts distributed across individual neuroanatomical brain regions for each study. Right panels present UpSet matrices and corresponding horizontal bar charts delineating the intersection sizes and structural distribution of key clinical variables, including age epochs (bins spanning from the 20s to the 90s), donor sex (male, m; female, f), and overall neuropathological diagnosis (Control, CTR; Alzheimer’s disease, AD; or unknown status). Embedded central bar charts indicate the discrete distribution of available tissue donors profiled across varying numbers of distinct brain regions per individual.

**Fig. S3.**
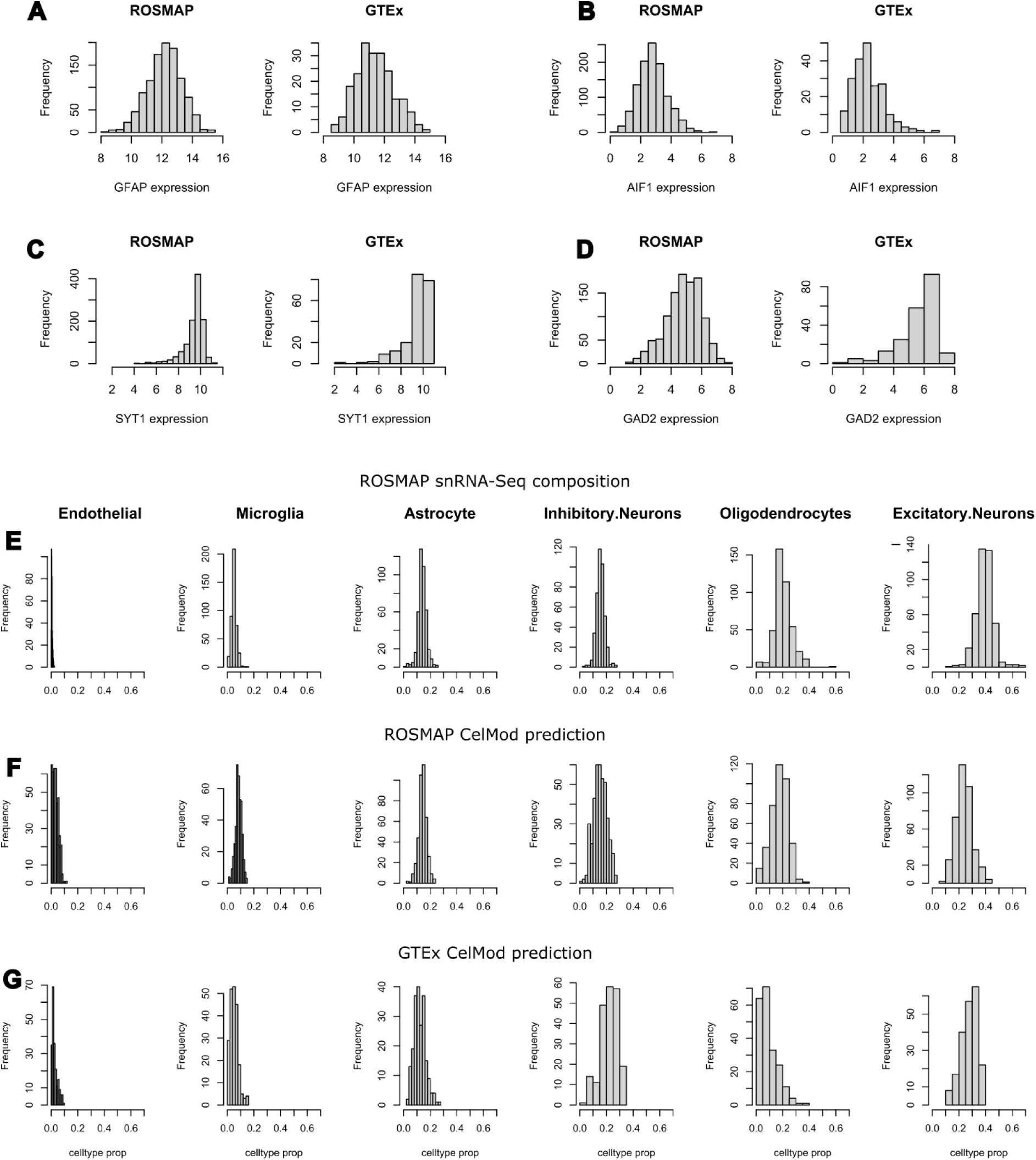
Evaluation of CelMod cellular deconvolution feasibility **(A to D)** Frequency distribution histograms displaying normalized gene expression levels for canonical central nervous system (CNS) lineage markers—*GFAP* for astrocytes (A), *AIF1* for microglia (B), *SYT1* for pan-neurons (C), and *GAD2* for GABAergic inhibitory neurons (D)—compared between the ROSMAP and GTEx DLPFC bulk transcriptomic datasets. Count matrices were uniformly processed and normalized using identical computational pipelines to confirm cross-study expression alignment. **(E)** Histograms highlighting the ground-truth distribution of relative proportions for major CNS cell types derived from the ROSMAP single-nucleus RNA sequencing (snRNA-Seq) reference cohort. Cellular lineages are organized left-to-right by increasing baseline abundance (from endothelial cells to excitatory neurons). **(F and G)** Histograms illustrating the relative cellular proportions predicted computationally via the CelMod algorithm within the bulk transcriptomic profiles of the ROSMAP (F) and GTEx (G) cohorts, validating a stable and comparable deconvolution architecture across both aging and neurodegenerative tissue cohorts.

**Fig. S4.**
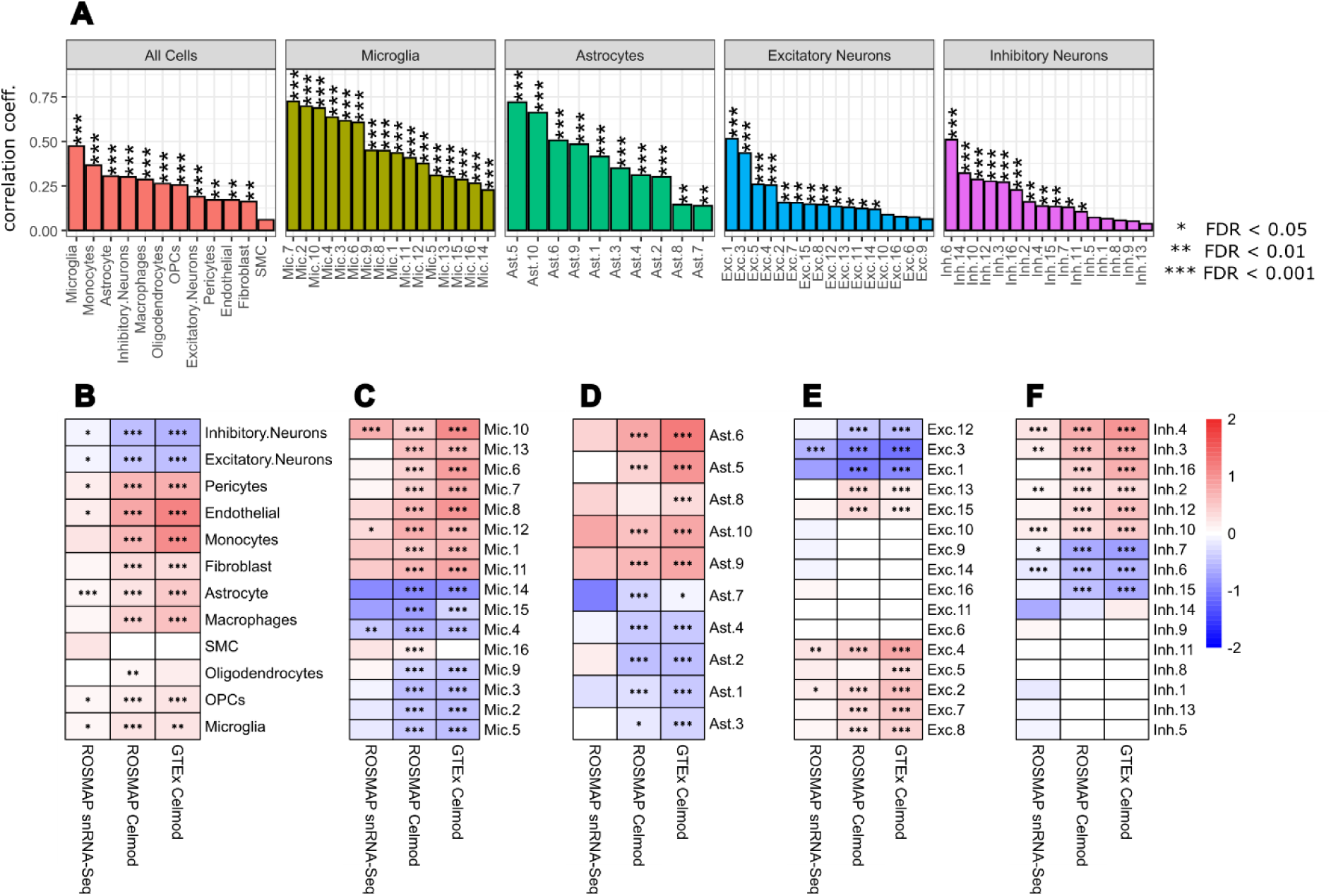
Comparison of the cellular composition of the immune-type DLPFC in snRNA-Seq and CelMod datasets (ROSMAP and GTEx) **(A**) Performance evaluation of CelMod-derived cellular and subpopulation proportions compared directly against ground-truth single-nucleus RNA sequencing (snRNA-Seq) measurements from matched tissue donors (n = 421). Bar heights indicate Spearman correlation between snRNA-seq and CelMod predicted proportions, over the test set of samples (Methods). Asterisks indicate statistical significance: *FDR < 0.05, **FDR < 0.01, ***FDR < 0.001. **(B to F)** Multi-dataset heatmaps illustrating shifts in the relative abundance of major CNS lineages (B), microglial subclusters (C), astrocyte subclusters (D), excitatory neuron subclusters (E), and inhibitory neuron subclusters (F) comparing immune-type to other-type samples. Cell values reflect log-transformed effect sizes derived from quasi-binomial regression models adjusted for study-specific clinical and demographic covariates across three distinct platforms: ROSMAP snRNA-Seq, ROSMAP bulk-level CelMod predictions, and GTEx bulk-level CelMod predictions.

**Fig. S5.**
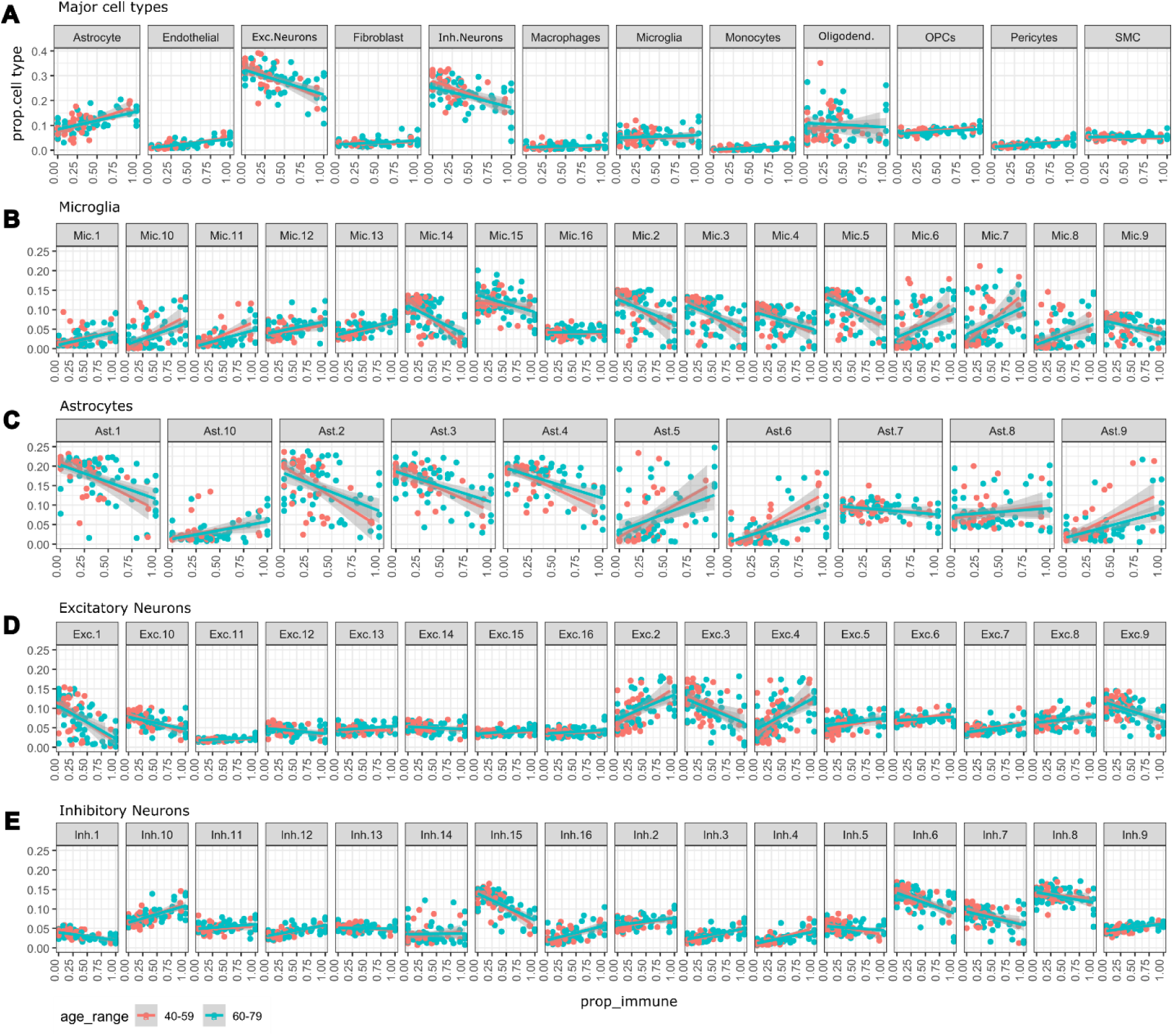
Changes in cellular composition with multi-region immune-type burden in GTEx DLPFC **(A to E)** Multi-panel scatter plots and fitted linear regression models mapping the estimated abundance of major cell types (A), microglial subpopulations (B), astrocyte subpopulations (C), excitatory neuron subpopulations (D), and inhibitory neuron subpopulations (E) as a function of brain prop_immune scores within the GTEx frontal cortex (FCx). Data points and regression trajectories are stratified and color-coded by donor age epochs, separating middle-aged (40–59 years, red) from elderly (60–79 years, cyan) cohorts. Fine-grained cellular subpopulations correspond to the high-resolution definitions established in Green et al., 2024 (20). Comprehensive statistical metrics and regression parameters are provided in Table S1.

